# Human T follicular helper clones seed the germinal center-resident regulatory pool

**DOI:** 10.1101/2022.10.26.513910

**Authors:** Carole Le Coz, Derek A. Oldridge, Ramin S. Herati, Nina De Luna, James Garifallou, Emylette Cruz Cabrera, Jonathan P Belman, Dana Pueschl, Luisa V. Silva, Ainsley V. C. Knox, Samuel Yoon, Karen B. Zur, Steven D. Handler, Hakon Hakonarson, E. John Wherry, Michael Gonzalez, Neil Romberg

## Abstract

How FOXP3^+^ T follicular regulatory (Tfr) cells simultaneously steer antibody formation toward microbe/vaccine recognition and away from self-reactivity remains unsettled. To explore human Tfr cell provenance, function and location heterogeneity, we used paired *TCRVA*/*TCRVB* sequencing to distinguish tonsillar Tfr cells clonally related to natural Tregs (nTfr) from those likely induced from Tfh cells (iTfr). The proteins iTfr and nTfr cells differentially expressed were utilized to pinpoint their *in situ* locations via multi-plex microscopy and establish divergent functional roles. *In-silico* and tonsil organoid tracking models corroborated the existence of separate Treg-to-nTfr and Tfh-to-iTfr developmental trajectories. In total, we have identified human iTfr cells as a distinct CD38-expressing, GC-resident, Tfh-descended subset that gains suppressive function while retaining capacities for B-cell help whereas CD38^-^ nTfr cells are elite suppressors primarily localized to follicular mantles. Interventions differentially targeting Tfr subsets may provide therapeutic opportunities to boost immunity or more precisely treat autoimmune diseases.

**One sentence summary:** Human tonsillar Tfr clones descend from either Treg or Tfh lineages and provenance predicts their TCR repertoires, locations and functional characteristics.

## INTRODUCTION

Germinal centers (GCs) are formed within secondary lymphoid tissues to orchestrate dynamic interactions between T follicular helper (Tfh) cells and GC B cells (*1*). GCs are remotely controlled by negative feedback from their chief product, systemically circulating affinity-maturated antibodies (*2*), and locally governed by FOXP3^+^ T follicular regulatory (Tfr) cells (*3*). Various Tfr cell-deficient mouse models demonstrate increased production of autoantibodies (*4,5*) and diminished vaccine responses (*6–9*). These findings suggest murine Tfr cells are functionally heterogeneous and express a mixture of T cell receptors (TCRs), some recognizing self, others recognizing foreign antigens.

The provenance(s) of Tfr cells is unsettled. Mouse Tfr cells were originally described as descending solely from thymically derived FOXP3^+^ T regulatory cells (Tregs) (*7–9*) but more recently, three animal studies have challenged this dogma. One demonstrated that vaccines containing incomplete Freund’s adjuvant could spur mouse FOXP3^-^ naïve CD4^+^ T cells to differentiate into vaccine-specific Tfr cells (*10*). A second study utilized longitudinal intravital microscopy to identify accumulations of Tfh-descended FOXP3-expressing T cells in murine GCs two to three weeks after immunization with an Alum-adjuvanted vaccine (*11*). Finally, *TCRA* sequence sharing between *TCRB*-transgenic murine CD4^+^ subsets indicated Tfr cells are most clonally related to Tregs and to a slightly lesser degree, Tfh cells (*12*). Notably, to collect sufficient *TCRA* mRNA for analysis, this group pooled material from multiple draining lymph nodes which likely diluted clonal relationships between cells from the same GC.

Although their function is likely important to immunologic health and their dysfunction a contributor to various disease states, few studies have assessed the biologic roles of human Tfr cells (*13–17*) and none have addressed their provenance(s) or intermediate developmental stages within tissues. Successful human Tfr cell investigations have been impeded by a lack of experimental tools to dynamically study human GC responses. Herein, we describe a series of novel approaches for studying human tonsillar Tfr cells including use of immunized tonsillar organoids as model systems to *in-vitro* track Treg and Tfh cell contributions to the Tfr pool. Further, we demonstrate how an interlocking suite of single cell technologies can clonally distinguish Tfr cells induced *in vivo* from Tfh lineage cells (iTfr) from those “naturally” derived from Tregs (nTfr). Once identified, we detail key genes and proteins distinguishing these two ancestrally distinct populations, including CD38, which is a reliable tonsillar iTfr cell biomarker. Using CD38 as a handle, we catalogued the precise *in situ* locations of iTfr cells in human tonsil tissues using co-detection by indexing (CODEX) multiplex microscopy and sort-separated iTfr from nTfr cells for downstream functional assessments. In total, we have identified heretofore unappreciated clonal, transcriptional, functional and positional heterogeneity in the human Tfr pool that reflect distinct descendant-ancestor relationships with both the Treg and Tfh cell lineages.

## RESULTS

### Differential CD25/CXCR5/PD1 expression distinguish tonsillar CD4^+^ T-cell subset transcriptomes

We isolated CD4^+^ T cells from excised pediatric tonsils and used CD25 expression to separate CD25^hi^ regulatory subsets from CD25^-^ Tfh cells, a strategy employed frequently in published mouse and human studies (Fig. 1A and fig. S1A) (*18*). Among the CD25^-^ population CXCR5 and PD1 staining defined Tfh cells with high purity as assessed separately with intracellular BCL6 and FOXP3 staining. BCL6 expression was universally detected in the CD25^-^CXCR5^+^PD1^hi^ Tfh gate with only a trivial frequency (<0.1%) of FOXP3 expressing cells identified (fig. S1B). Although a CD25^-^ Tfr subset has been described in mice and humans (*19,20*), less than 10% of the tonsillar CXCR5^+^FOXP3^+^ population analyzed lacked CD25 expression and since none of these highly expressed PD1, they could not be mistaken for Tfh cells (fig. S1C).

**Figure 1.**
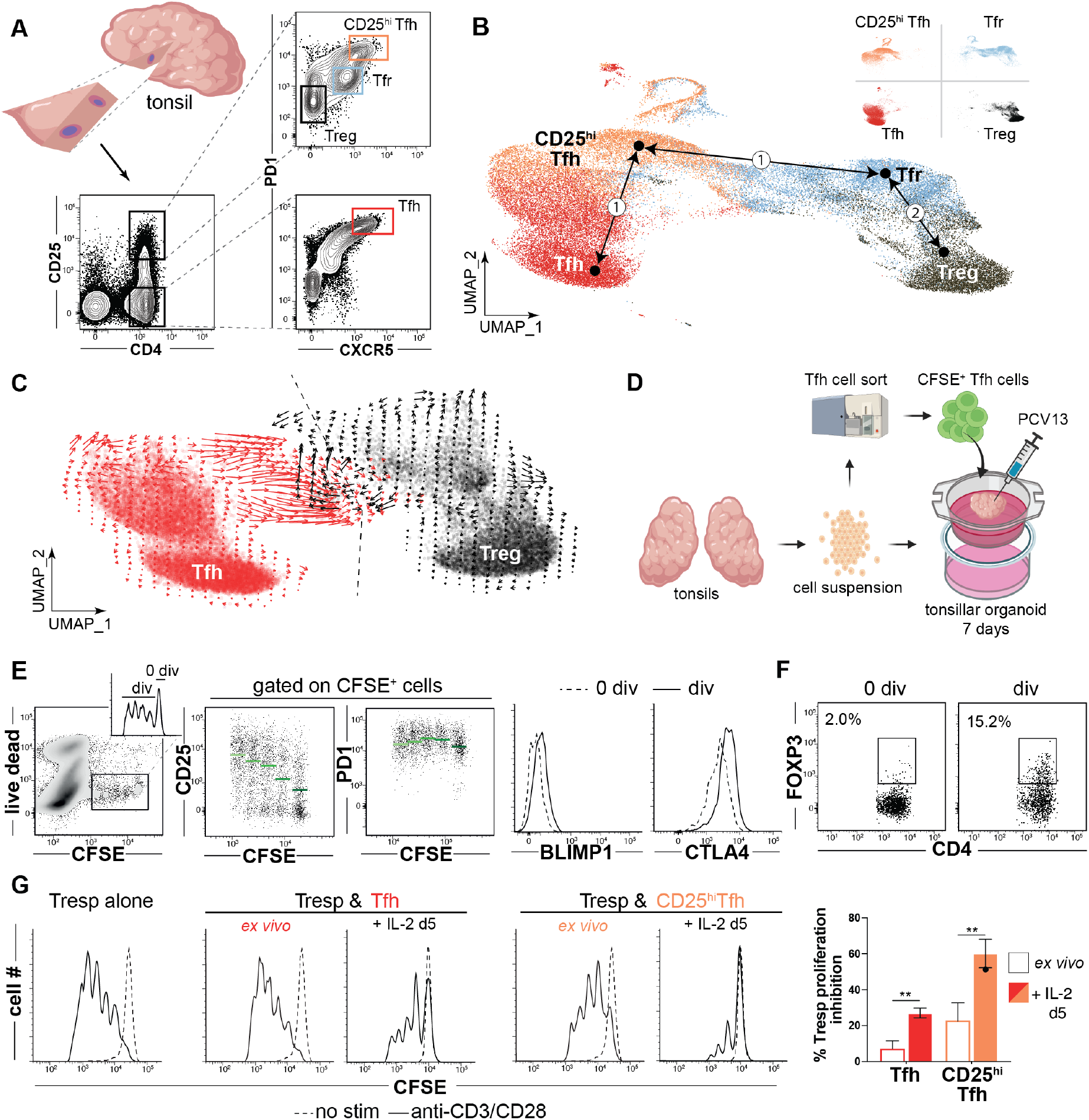
*In silico* and *in vitro* models predict Tfh and Treg cells each contribute to the Tfr pool. (**A**) A strategy to sort Tfh, CD25^hi^Tfh, Tfr, and Treg cell subsets from a small tonsil wedge is depicted. (**B**) Dimensionally reduced single cell RNA-sequencing of TC174 T cell subsets are displayed as a UMAP. Pseudotime cell ordering projections (black arrows) identify two possible Tfr developmental arcs, one to CD25^hi^Tfh and Tfh cells (1), the second to Tregs (2). (**C**) RNA velocities of Tfh and Treg cells from TC174 are displayed. An arbitrary dashed centerline denotes convergence. (**D**) A strategy to track within vaccinated tonsillar organoids is depicted. (**E**) CD25, PD1, BLIMP1, CTLA4 and (**F**) FOXP3 expression by divided (div) and non-divided (0 div) CFSE stained Tfh cells are displayed 7 days after PCV13 vaccination. Green horizontal bars indicate mean values (**G**) Histograms record inhibition of T responder (Tresp, CD45RO^-^CD25^-^ CD4^+^ T cells) cell proliferation in co-cultures with Tfh cells or CD25^hi^Tfh cells before or after five days of recombinant IL2 treatment. **, *P*<0.01 by Mann-Whitney U tests.

Within the CD25^hi^ subset were classic CXCR5^-^PD1^lo^ Tregs, which expressed FOXP3, and two CXCR5-expressing follicular populations. One CXCR5^+^ subset, which stained PD1 intermediate, corresponded to Tfr cells since it expressed higher FOXP3 than Tregs and could suppress T responder (Tresp) cell proliferation *in vitro* (fig. S1D). The other CXCR5^+^ population was PD1^hi^ and secreted IL-10, but did not express FOXP3 (fig. S1A). Although a similar CD25^hi^CXCR5^+^PD1^hi^, IL-10 secreting tonsillar subset was attributed suppressive function by another group (*21*), in our hands these cells were not effective at inhibiting T responder (Tresp) proliferation *in vitro* (fig. S1D) (*22*). For this reason, we descriptively call this population “CD25^hi^Tfh”.

To explore transcriptional relationships between tonsillar CD4^+^ T cells we employed the above strategy to sort live Treg, Tfr, CD25^hi^Tfh and Tfh subsets from two immunocompetent pediatric tonsil donors, TC174 and TC341. Rather than utilizing entire tonsils or tonsil pairs, fewer cells (20,000 per subset) were collected from small tonsil wedge dissections to selectively reduce the number of GCs sampled and to maximize clonal overlaps between antigen-expanded lineages (Fig. 1A). Subset-specific single cell RNA-sequencing (sc-RNAseq) libraries were created from sorted cells with an average of 10,372 cells recovered per library (range 8,512-15,598 cells, table S1). Sequenced libraries were dimensionally reduced to form a single two-dimensional Uniform Manifold Approximation and Projection (UMAP) for each tonsil donor (Fig. 1B and fig. S2). In both UMAPs Treg and Tfh cells localized to distant, non-overlapping spaces while Tfr cells and CD25^hi^Tfh cells filled the intervening space. As expected from our sorting strategy, *CXCR5* and *PDCD1* (PD1) transcripts were highest in CD25^hi^Tfh and Tfh cells. *IL2RA* (CD25) transcripts were lowest in Tfh cells (fig. S3). *FOXP3* transcripts were predictably enriched in Treg and Tfr cells. *PRDM1* (BLIMP1) and *CTLA4* expression were greatest in Tfr, but also high in CD25^hi^Tfh cells. Thus, sorting on variable CD25, CXCR5, and PD1 expression defines four transcriptionally distinct subsets, two that expressed FOXP3 and two that did not. This strategy efficiently captures the large majority of the Tfr cell population while effectively preventing cross contamination between Tfh and Tfr cells.

### *In-silico* and *in-vitro* models predict Treg and Tfh/CD25^hi^Tfh lineages each contribute to the Tfr pool

To explore the competing hypotheses that Tfr cells descend from Tregs or from Tfh cells, we set Treg, Tfr, CD25^hi^Tfh and Tfh cell transcriptomes as conceptual differentiation start/end points and used pseudotime analysis (*23*) to infer developmental trajectories (but not trajectory directions) based on progressive gene expression states. Two Tfr developmental arcs were generated; one connected Tfr cells directly to Tregs and the other connected Tfr to Tfh cells by passing through CD25^hi^Tfh cells (Fig. 1B). To deduce trajectory directions, we compared the relative abundance of nascent (unspliced) and mature (spliced) mRNA of single Tfh and Treg cells (i.e. RNA velocity). Although transcriptionally distinct, Treg and Tfh RNA velocities converged along a centerline that ran directly through Tfr transcriptional space (Fig. 1C). Hence, results from transcriptionally-based, *in-silico* developmental models are consistent with two distinct ancestor-descendant relationships for Tfr cells, one shared with Tregs and the other shared with CD25^hi^Tfh and/or Tfh cells.

To test pseudotime and RNA velocity predictions *in vitr*o, tonsillar organoids, which generate vaccine-specific immune responses within characteristic GC light and dark zones (*24,25*), were utilized to dynamically track Treg and Tfh lineage cells. In separate experiments, 1×10^4^ sorted CellTrace™ Violet (CTV)-stained Tregs or 4×10^4^ sorted carboxyfluorescein succinimidyl ester (CFSE)-stained Tfh cells were incorporated into autologous tonsillar organoids. Organoids were subsequently vaccinated with a human pneumococcal diphtheria toxin-conjugated vaccine adjuvanted with Alum (PCV13, illustrated in Fig. 1D and fig. S4A). After seven to eight days in culture, organoid cells were mechanically dissociated, resuspended and intravitally stained cells re-identified by flow cytometry. Although most CellTrace™ Violet-stained Tregs did not divide in organoids, the 1-2% that did divide maintained FOXP3 expression and differentially upregulated CXCR5 4-7 fold, a change consistent with Tfr differentiation (fig. S4, B and C). The frequency of divided CFSE-stained Tfh cells was greater (52%), with some cells demonstrating ≥ 3 divisions. With each division, CD25 mean fluorescence intensities (MFIs) increased whereas PD-1 expression remained stable (Fig 1E and fig. S4D). Also, compared with undivided cells, divided CFSE-stained Tfh cells upregulated key regulatory proteins including BLIMP1, CTLA4, and FOXP3 (Fig. 1E, F and fig. S4D). Similar changes were observed by culturing either tonsillar Tfh cells or CD25^hi^Tfh cells with recombinant IL-2 (rIL-2) and antiCD3/CD28. After five days of rIL2 treatment, Tfh cells slowly lost BCL6 expression and upregulated CD25^hi^Tfh-associated proteins including BLIMP1, CD25, CTLA4 and Ki67 but not FOXP3 (fig. S5A and B). In contrast, CD25^hi^Tfh cells treated with rIL2 for five days began expressing FOXP3 (fig. S5B) and were significantly better at suppressing Tresp cell proliferation than *ex vivo* CD25^hi^Tfh (p<0.01; Fig. 1G). Hence, *in-vitro* models suggest IL-2 responsive tonsillar Tfh cells pass through a proliferative CD25^hi^Tfh intermediate stage to seed the Tfr pool.

### Tfr clones are related to either Treg or Tfh/CD25^hi^Tfh lineages, but not both

To determine what clonal relationship(s) exist between human tonsillar CD4^+^ T cell subsets *in vivo*, paired *TCRB* and *TCRA* transcripts were analyzed from each of TC174 and TC341’s respective scRNA-seq libraries. An average of 7,873 cells per library (range 6,484 to 9,596 cells, table S1) were recovered with both a *TCRB* and *TCRA* nucleotide sequence. Although no consistent differences in subset *V*(*D*)*J* gene usage patterns (fig. S6 and S7) and predicted CDR3 amino acid length distributions were appreciated (fig. S8), Treg clonal diversity was considerably greater than Tfr, CD25^+^Tfh and Tfh diversity by several measures (Shannon’s diversity index, inverse Simpson index, Chao1 and abundance-based coverage estimator (ACE); Fig. 2A). These measures likely reflect the higher proportion of non-expanded naïve cells in the Treg pool relative to other studied subsets (fig. S9). For instance, 96.2±0.3% of Treg clones were comprised of a single cell, a frequency considerably higher than in Tfh (85±1.0%), CD25^hi^Tfh (76.5±1.5%), and Tfr (78±2.0%) subsets (Fig. 2B and fig. S10A). Similarly, the top ten largest Treg clones (which numbered 16.5±4.4 clones due to ranking ties) were comprised of only 10.1±5.5 cells, whereas Tfh, CD25^hi^Tfh, and Tfr top ten clones were bigger, containing 24.1±7.5 cells, 46±14.1 cells, 32.5±7.7 cells, respectively. These data suggest follicular subsets like Tfh, CD25^hi^Tfh, and Tfr are more clonally-expanded than Tregs.

**Figure 2.**
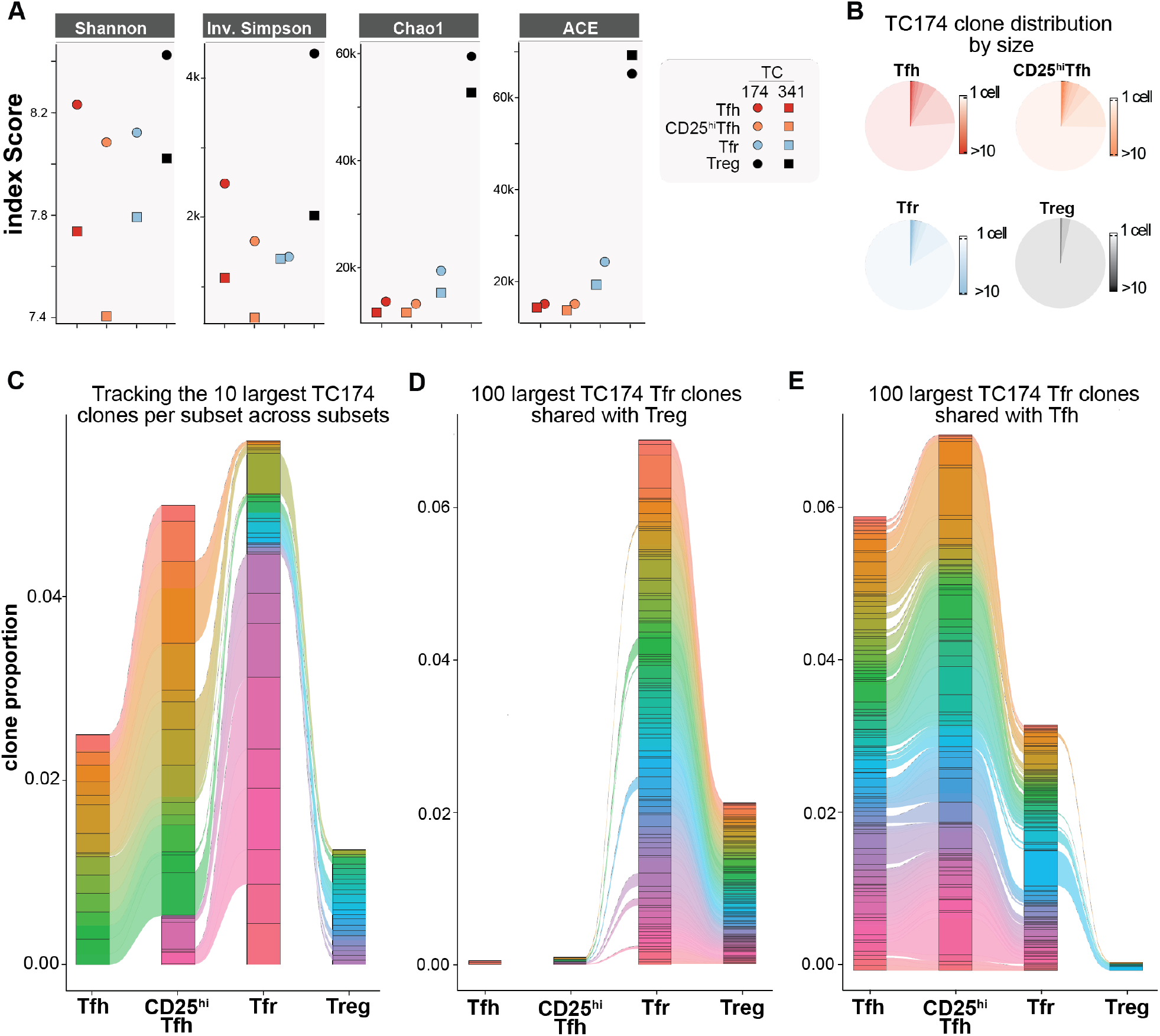
The tonsillar Tfr pool shares clones with clonally diverse Tregs and clonally expanded Tfh/CD25^hi^Tfh cells. (**A**) *TCRA/TCRB* repertoire diversities of Tfh, CD25^hi^Tfh, Tfr and Treg subsets from TC174 and TC341 are measured using Shannon, Inverse Simpson, Chao and ACE indices. (**B**) Pie charts indicate subset-specific clone size distributions (cells per clone) for TC174 (**C**) Tracking the top ten largest clones per TC174 subset across other subsets, (**D**) tracking the top 100 largest clones shared between TC174 Tfr and Tfh cells and (**E**) between TC174 Tfr and Tregs across other subsets. More than 10 clones or 100 clones per subset are sometimes displayed to account for ranking ties and clone sharing across subsets.

In mice, the TCRαβ repertoires of thymically-derived Tregs and FOXP3^-^ T helper cells do not overlap (*12,26*). Like published murine data, we found clone sharing between Treg and Tfh cells from the same tonsil donor to be minimal (0.3±0.1%), with no overlap among either donor subsets’ top ten largest clones (Fig. 2C, fig. S10B and fig. S11). In contrast, tonsillar Tfh and CD25^hi^Tfh cells from the same donor shared many clones (11.5±0.5%), including most (93.2±1.9%) of the same top ten largest clones reinforcing Tfh and CD25^hi^ cell lineage affiliation. As predicted by the mouse literature (*12*), many (29±21%) top ten Treg clones were shared with Tfr cells but none of these were among the largest Tfr cell clones. Instead, some of the largest Tfr clones were shared with either Tfh or CD25^hi^Tfh lineage cells. Moreover, none of the top ten largest clones shared between Tfr and Treg cells overlapped with the top ten largest clones shared between Tfr and Tfh/CD25^hi^Tfh lineage cells (Fig. 2C and fig. S10B), indicating divergent Tfr ancestry.

To determine how comprehensively the Tfr pool could be divided by its clonal relationships with Treg and Tfh/CD25^hi^Tfh lineage cells, we expanded our analysis beyond the top ten to include all Tfr clones (8,207±16 clones). Through this more exhaustive approach we identified a total of 150±85 clones shared between Tfr and Tregs (Fig. 2D, fig. S10C and fig. S10E) and 318±58 Tfr clones shared between Tfr and Tfh cells (Fig. 2E, fig. S10D and fig. S10E). Importantly, these three subsets only held 5.5±0.5 clones in common reinforcing that Tfh and Treg cells separately contribute to the Tfr pool. Also notable was the high degree of clone sharing between Tfr, CD25^hi^Tfh and Tfh cells. 50.4±8% of the clones shared by Tfr and Tfh cells were also shared between Tfr and CD25^hi^Tfh cells consistent with a Tfh to CD25^hi^Tfh to Tfr developmental arc. In contrast, only 7.6±1.7% of Tfr clones shared with Tregs were also shared with CD25^hi^Tfh cells. Hence, corroborating *in-silico* predictions and *in-vitro* lineage tracing, the majority of shared tonsillar Tfr clones overlap either Treg lineage or Tfh/CD25^hi^ lineage cells but importantly not both populations. These observations provide *in-vivo* evidence that Tregs and Tfh clones seed the human tonsillar Tfr pool separately.

### Stringent clonal relationships distinguish Tfr cells with divergent ancestries

Although sort purity was uniformly high (fig. S12), we could not exclude the possibility that some observed clonal overlaps between tonsillar CD4^+^ T cells subsets were the result of unintentional contamination. To reduce experimental noise we performed a second, more strict analysis which defined a “stringent clone” as ≥ 2 cells in the same donor subset that expressed identical paired *TCRA* and *TCRB* nucleotide sequences. This more rigid definition further minimized the clonal overlap between Tregs and Tfh cells, leaving only one stringent clone in the combined TC174 and TC341 dataset (Fig. 3A). Other clonal relationships were strengthened by stringent analysis including Tfr and Treg cells (42 combined stringent clones), Tfr and Tfh cells (73 combined stringent clones), and Tfh cells and CD25^hi^Tfh cells (603 combined stringent clones).

**Figure 3.**
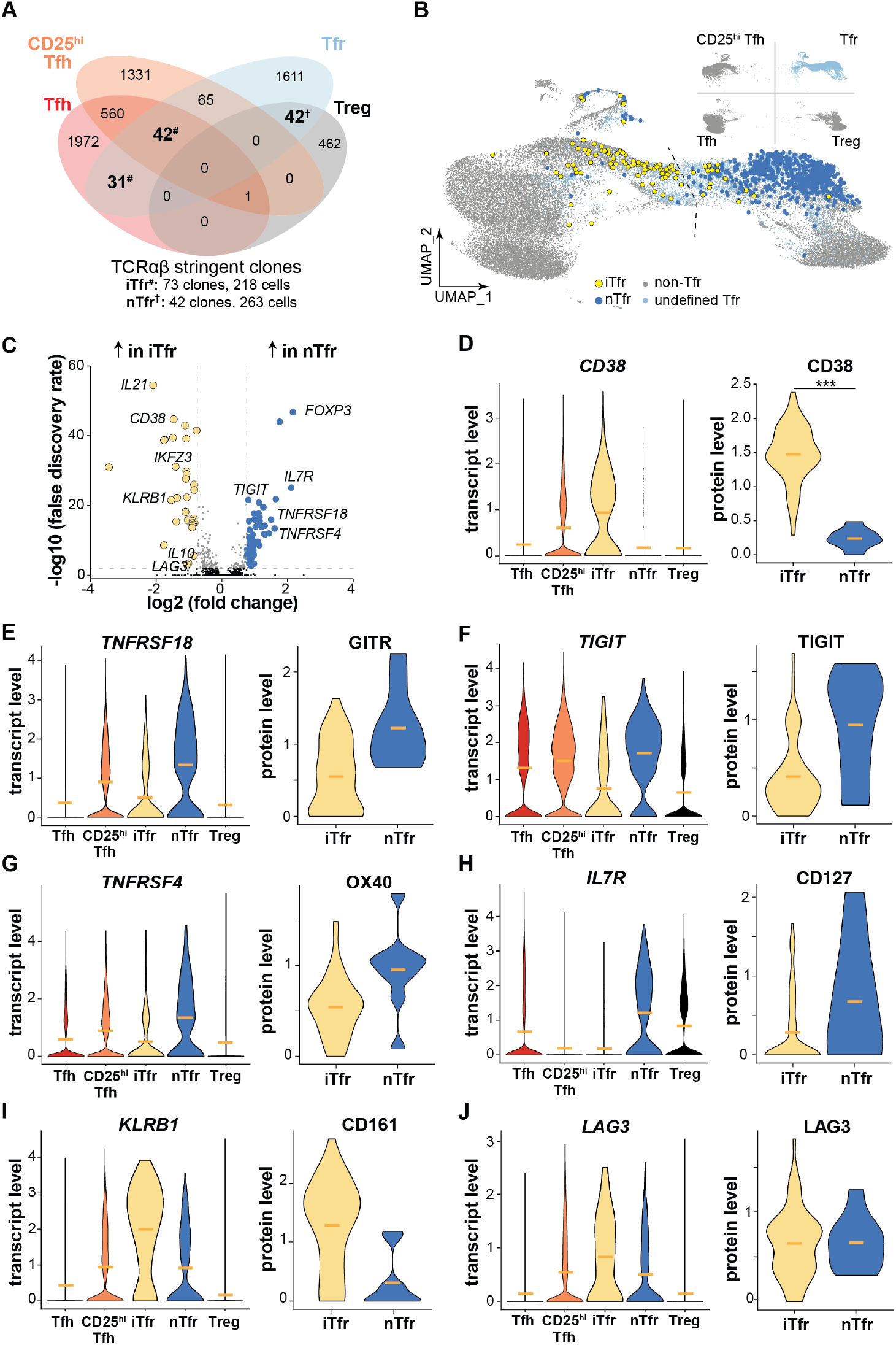
Transcriptomes of stringently defined iTfr and nTfr cells diverge and are predicted by variable *CD38* expression. (**A**) Venn diagram displays overlapping TC174 stringent clones, defined as a clone with ≥2 cells. #, iTfr stringent clones; †, nTfr stringent clones. (**B**) Positions of stringent TC174 iTfr and nTr cells are displayed overlying a UMAP plot. (**C**) A volcano plot shows genes differentially expressed between stringently defined iTfr and nTfr cells pooled from TC174 and TC341. (**D-J**) Violin plots display differential expression of pooled transcripts and corresponding cell surface protein expression on TC341 subsets. Horizontal bars indicate mean values. ***, *P*<0.001 by Wilcoxon rank sum test with Bonferroni correction.

To group Tfr cells by their clonal relationships with other subsets, Tfr cells that shared stringent clones with Tregs, but not Tfh cells, were designated “nTfr” cells. The “n” references the natural Treg->Tfr differentiation pathway originally described in mice *(7– 9*). Separately we designated Tfr cells that shared stringent clones with Tfh cells, but not Tregs, as “iTfr” cells (so named because they were likely induced from Tfh lineage cells). On tonsil donor-specific transcriptomic UMAPs, most iTfr cells localized closer to Tfh transcriptional space, whereas nTfr cells localized nearer to Tregs (Fig. 3B and fig. S13). Furthermore, few Tfr cells of either kind crossed the centerline previously determined through RNA-velocity analysis of TC174 cells (Fig. 3B). These data suggest stringent clonal relationships between tonsillar T helper subsets capture lineage-specific transcriptional differences.

### Surface CD38 expression effectively distinguishes tonsillar iTfr and nTfr cells

To explore transcriptional differences between iTfr and nTfr cells, we pooled scRNA-seq data from the clonally identified iTfr cells (n=218) and nTfr cells (n=263) from both tonsils and found 86 differentially expressed genes (DEGs, Fig. 3C). DEGs upregulated by nTfr cells connected closely to Treg biology and included *FOXP3, ENTPD1* (CD39), *TIGIT, TNFRSF4* (OX40), and *TNFRSF18* (GITR). Notably *IKZF2* (HELIOS), once considered a practical biomarker to distinguish natural, thymically-derived Tregs from induced counterparts, was uniquely expressed by nTfr cells (*27*). DEGs upregulated by iTfr cells encoded the follicular cytokine IL21, the regulatory cytokine IL10, AIOLOS (*IKZF3*), and LAG3 (Fig. 3C). iTfr cells also uniquely expressed *CD38* and lacked nearly all *IL7R* transcripts. Of all DEGs encoding cell surface proteins, *CD38* was one of the most promising candidate biomarkers to potentially distinguish iTfr from nTfr cells (Fig. 3D). Additionally, DNA-barcoded antibodies targeting 139 TC341 Tfr cell surface molecules identified CD38, over GITR, TIGIT, OX40, CD127, CD161 and LAG3, to be the protein most significantly and consistently upregulated by iTfr cells relative to nTfr cells (p<0.001) (Fig. 3D-J).

To further validate CD38 as a potential iTfr biomarker, a flow cytometric survey of Tfr cells was performed on five to seven additional, unrelated pediatric tonsil donors. Across tonsil donors CD38 staining was consistently bimodal with approximately 35% of Tfr cells expressing it and 65% not (Fig. 4, A and B), a proportion that matched the ratio of clonally defined nTfr to iTfr cells (Fig. 3A). CD38^-^Tfr cells also expressed more CD39, CD25, CTLA4, CD127, GITR, HELIOS, TIGIT, and FOXP3 than CD38^+^Tfr cells, consistent with clonally defined nTfr transcriptomes (Fig. 4C-G and fig. S14). CD38^+^Tfr cells were uniformly AILOS^+^LAG3^+^CD127^-^ cells, a profile matching transcriptomes of clonally defined iTfr cells (Fig. 4C). ICOS, IL21, and Ki67 MFIs were also all significantly higher in CD38^+^Tfr cells (p<0.01, p<0.001 and p<0.0001, respectively; Fig. 4C) reflecting a likely developmental relationship with Tfh/CD25^hi^Tfh lineage cells. Hence, differential CD38 expression by tonsillar Tfr cells predicts broader, lineage-associated immunophenotypes.

**Figure 4.**
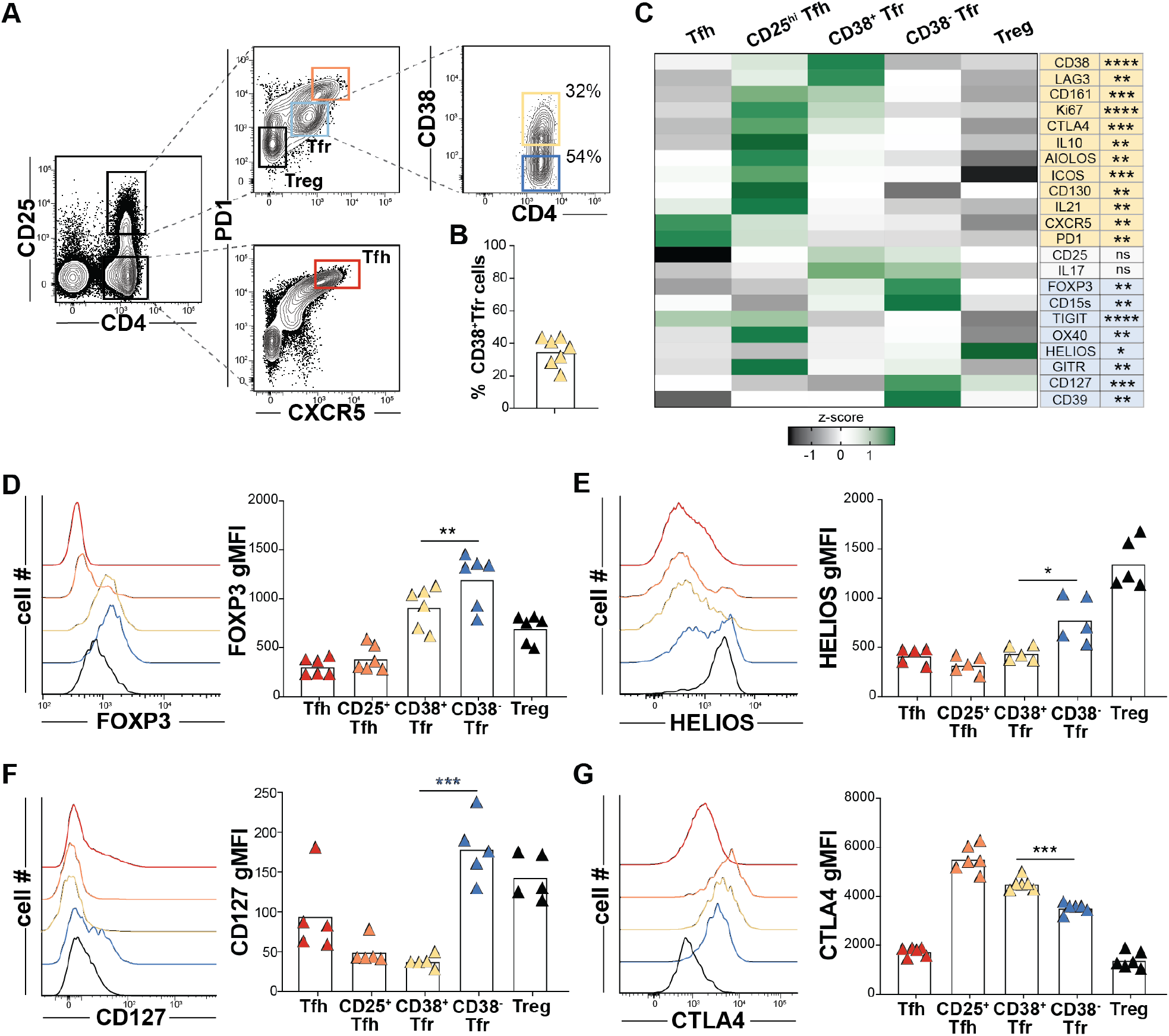
Variable CD38 expression predicts distinct and larger Tfr cell immunophenotypes. (**A**) Tfr expression of CD38 from a representative tonsil donor and (**B**) Tfr CD38 expression frequencies for 6 pediatric tonsil donors. (**C**) A heat map displays geometric mean fluorescence intensity (gMFI) z-scores associated by T-cell subsets from 6 donors. Proteins differentially upregulated in CD38^+^Tfr (yellow) and CD38^-^Tfr cells (blue) are shaded. (**D-G**) Histograms display gMFIs of indicated proteins by T subsets from a representative tonsil donor and bar graphs show gMFIs for 5-6 tonsil donors. *, *P*<0.05; **, *P*<0.01; ***, *P*<0.001; ****, *P*<0.0001 by Mann-Whitney U tests.

### CD38^+^>Tfr cells are regulatory cells specialized to provide B-cell help; CD38^-^ Tfr cells are elite suppressors

Although *FOXP3* transcripts and FOXP3 MFIs were significantly higher in nTfr cells and CD38^-^Tfr cells, respectively, CD38^+^Tfr cells clearly also expressed FOXP3 (Fig. 4D). In fact, FOXP3 MFIs in CD38^+^Tfr cells were as high as in *bona fide* tonsillar Tregs, suggesting a regulatory identity. Additionally, CD38^+^Tfr cells consistently demonstrated the highest CTLA4 MFIs and the greatest frequencies of IL10 secreting cells of any studied regulatory subset (Fig. 4, C and G and Fig. 5A). Indeed, sorted CD38^+^Tfr cells were as capable of inhibiting Tresp proliferation *in vitro* as tonsillar Tregs (Fig. 5B). More impressive suppressive function was demonstrated by CD38^-^Tfr cells which outperformed all other tested subsets in *in-vitro* assays (p<0.001 for both comparisons). Since CD38^-^ Tfr cells displayed lower CTLA4 MFIs (Fig. 4G) and less IL10 expression than CD38^+^Tfr cells (Fig. 5A), their elite suppressive function may derive from one or more other, previously described regulatory strategies (i.e. ectonucleotidase, IL-2 sink, GITR) (*28*).

**Figure 5.**
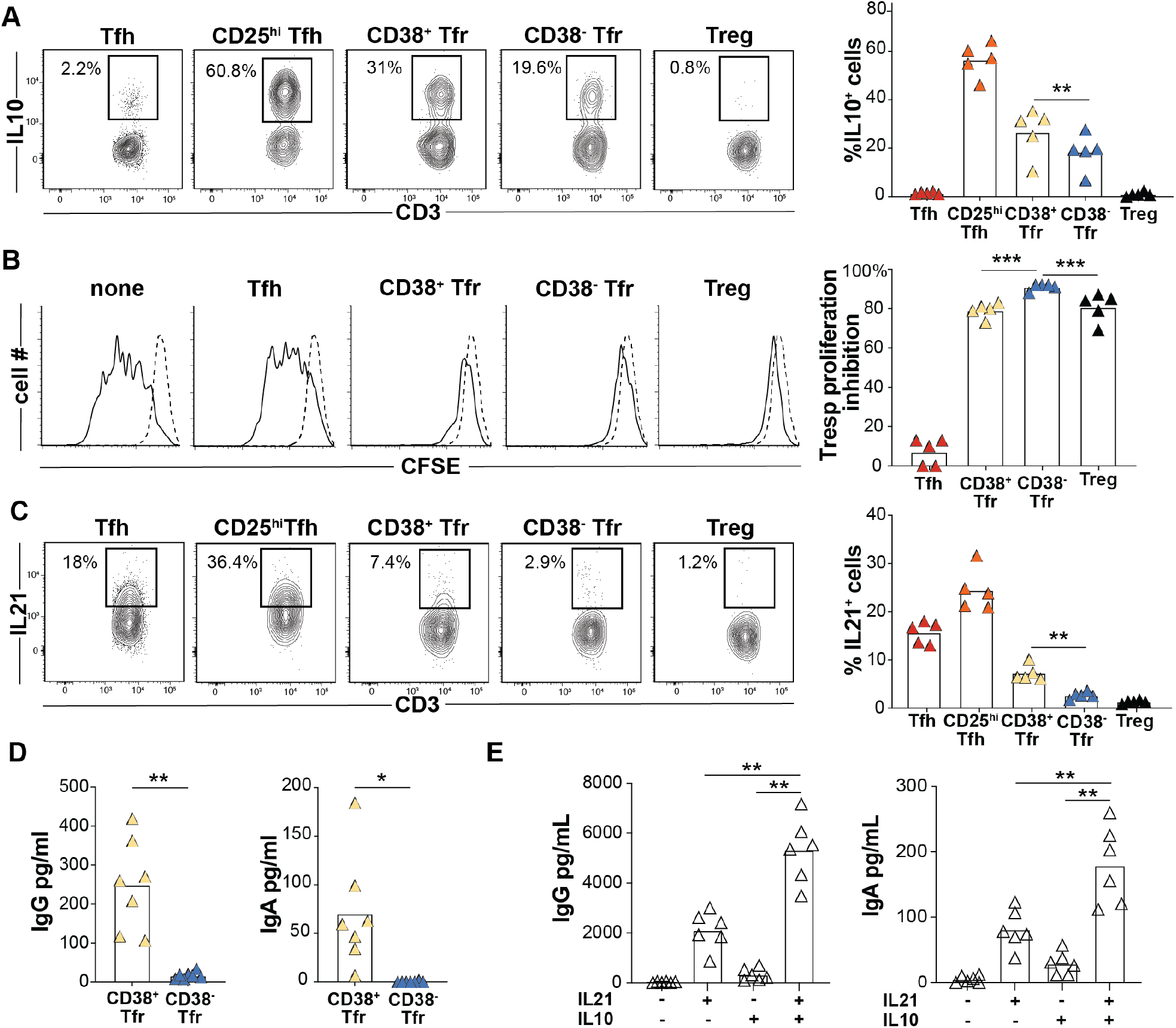
CD38^+^Tfr cells provide germinal center B-cell help. CD38^-^Tfr cells are elite suppressors. (**A**) Frequencies of PMA/ionomycin-activated T-subsets expressing IL10 are shown for a representative donor (left) and all available donors (n=5, right). (**B**) Day 4 Tresp cell proliferation with and without indicated T cell subsets from for a representative donor (left) and all available donors (n=5, right) (**C**) Frequencies of PMA/ionomycin-activated T-subsets expressing IL21 are shown for a representative donor (left) and all available donors (n=5, right). Day 7 IgG and IgA supernatant concentrations from GC B cells (**D**) co-cultured with either CD38^+^ or CD38^-^ Tfr cells stimulated with anti-CD3/CD28 beads (n=5) or (**E**) cultured alone with megaCD40L, recombinant IL21 and/or recombinant IL10 treatment (n=7). *, *P*<0.05; **, *P*<0.01; ***, *P*<0.001; by Mann-Whitney U tests.

Within GCs, Tfh cells induce B-cell class-switching, somatic hypermutation and plasmablast differentiation by secreting IL-21 (*29,30*). Since *IL21* was the most upregulated iTfr DEG (Fig. 3C), we compared the frequencies of IL21-secreting CD38^+^ and CD38^-^Tfr cells across several tonsil donors. Indeed, on average 2.7 times as many tonsillar CD38^+^Tfr cells secreted IL21 as CD38^-^Tfr cells (7.3% vs. 2.7%, p<0.01; Fig. 5C). Moreover, GC B cells co-cultured with autologous CD38^+^Tfr cells generated significantly higher supernatant IgG (p<0.01) and more IgA (p<0.05) than GC B cells co-cultured with CD38^-^Tfr cells (Fig. 5D). To determine if IL-10 or IL-21 secretion by CD38^+^Tfr cells could account for enhanced IgG and IgA secretion we cultured GC-B cells with and without recombinant IL-10 and/or IL-21. Although IL-10 had little effect by itself, GC B cells treated with a combination of IL-10 and IL-21 secreted significantly more IgG and IgA than those treated with IL-21 alone (p<0.01 for both comparisons; Fig.5 E). Hence, CD38^+^Tfr cells are regulatory cells that retain and refine their capacity for B-cell help through IL-21 and IL-10 secretion. CD38^-^Tfr cells are elite suppressors.

### CD38^+^Tfr cells reside inside GCs, CD38^-^Tfr cells localize to the follicular mantle

Although the follicular dendritic cell lattice attracts Tfh and Tfr cells by maintaining a CXCL13 gradient (*31,32*), not all CXCR5 expressing T helper cells are located within GCs. Prior analysis of human lymph nodes indicate most human Tfr cells accumulate within follicles but clearly outside GC borders (*20,33*). To identify the precise locations of Tfr cells in human tonsillar tissue we employed CODEX to sequentially stain and image a DAPI-stained donor tonsil with a panel of 23 oligo-conjugated antibodies. Antibody targets included those relevant to follicular T cell biology and lineage-defining antigens to identify B cell, T cell, follicular dendritic cell and epithelial populations (table S2). After image acquisition, stitching, registration, segmentation, normalization, and high-dimensional clustering by marker profiles to identify cell types, a k-nearest neighbors-based, machine-learning platform (*34*) was employed to assign all tonsillar cells – totaling more than 1.5 million– to one of ten cellular neighborhoods (CNs) (Fig. 6A). Each CN possessed a characteristic cell type composition profile (fig. S15A and B) (*34*). Tonsil follicles were comprised of three distinct neighborhoods, CN3, CN6, and CN7. CN3 and CN7 primarily contained either GC B cells or Tfh cells, respectively, and corresponded to GC dark and light zones (fig. S15 A-C). CN6 regions surrounded GCs and were comprised largely of naive B cells, a profile consistent with follicular mantles (fig. S15A-C). To determine the spatial distribution of Tfr cells, we searched cells in follicles (i.e. CN3, CN6 and CN7) for FOXP3 and CD4 co-expression (Fig. 6B). Of 1,206 Tfr cells identified in this manner, 1,164 localized to mantles (Fig. 6A and B). The remaining Tfr cells were dispersed among dark zones (n=25) and light zones (n=17; Fig. 6A and B). Tfr locations identified by CODEX were independently confirmed by analyzing five additional tonsil donors with conventional immunofluorescence confocal microscopy. Indeed, most Tfr cells (71%) identified by our secondary analysis were positioned outside of GCs (fig. S16C).

**Figure 6.**
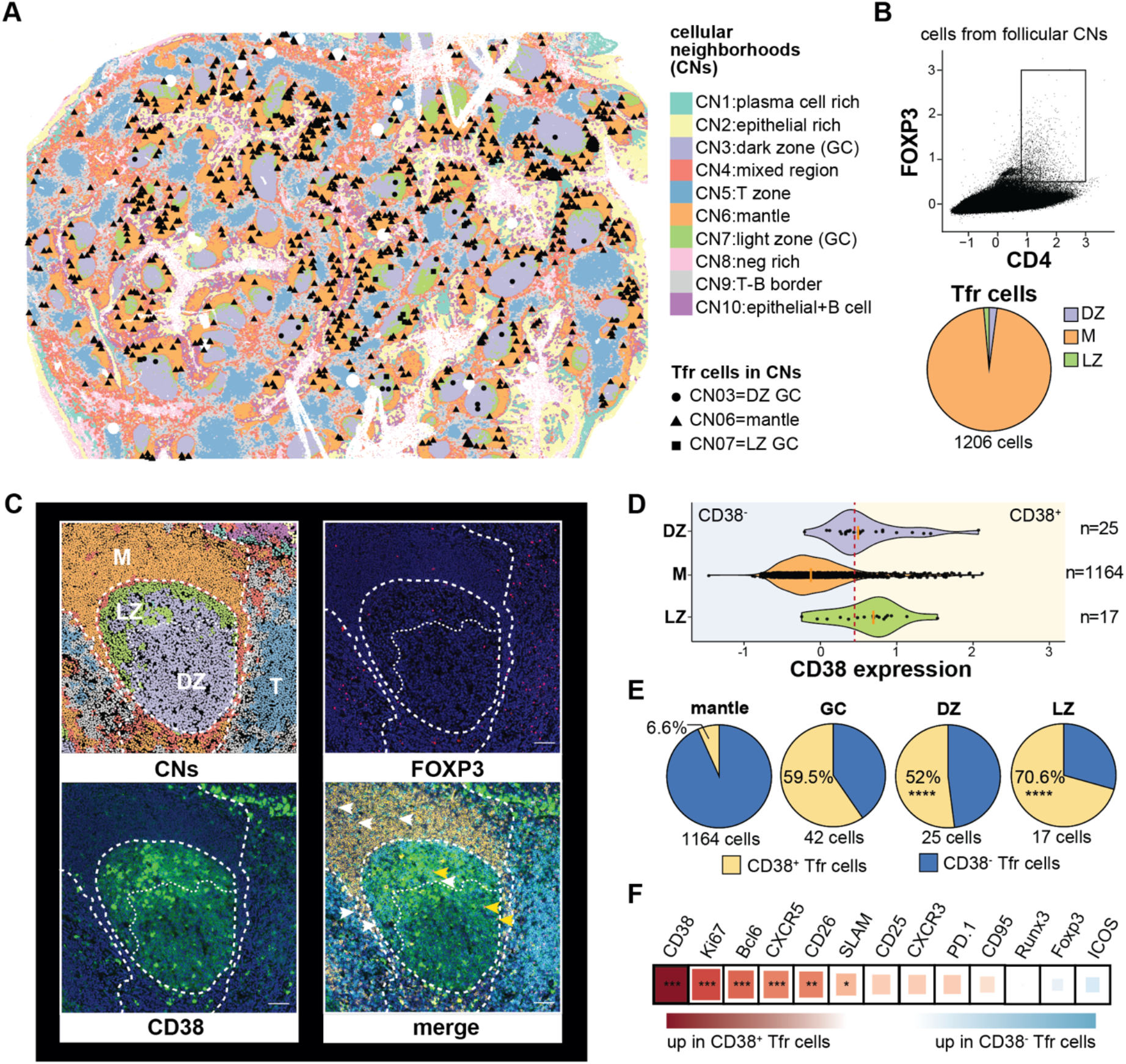
Multiplex imaging principally identifies CD38^+^Tfr cells within tonsil germinal centers. (**A**) A CODEX-stained human tonsil section is portioned into cell neighborhoods (CNs) and the locations of CD4^+^FOXP3^+^ cells in the dark zone (CN3), mantle (CN6), and light zone (CN7) are marked. (**B**) A CD4^+^FOXP3^+^ gate and the proportion of gated cells in each follicular CN is shown. (**C**) A representative tonsil GC with CNs, FOXP3 (pink) staining, CD38 (green) staining, and CD4 (cyan), IgD (orange), CD38, and FOXP3 overlaid. CD38^+^ (yellow arrow heads) and CD38^-^ Tfr cells (white arrows heads) positions are indicated (**D**) Violin plots indicate CD38 expression distribution on CD4^+^FOXP3^+^ cells from different CNs. A red dashed line indicates the expression threshold above which Tfr cells were considered CD38^+^. (**E**) Proportions of CD38 expressing subsets in indicated follicular CNs. (**F**) Differential protein expression by CD38^+^ and CD38^-^Tfr subsets. Square size and color indicate fold change difference and direction, respectively, by Spearman correlation R coefficient. *, *P*<0.05 **, *P*<0.01 ***, *P*<0.001, ****, *P*<0.0001 by unpaired Wilcoxon tests or Chi-squared test. Scale bar 200-pixel=64.5μm. DZ, dark zone; LZ, light zone; M, mantle; T, T cell zone.

CD38 is distributed across tonsillar mononuclear cells in a trimodal pattern (fig. S17). At the extremes of expression are naïve B cells, which do not substantively express CD38, whereas plasmablast/plasma cells and highly activated GC B cells stain CD38^bright^. Intermediate CD38 expressors include a proportion of GC B cells and tonsillar T cells (fig. S17). To assess follicular CD38 distribution *in situ*, we stained tonsil sections with anti-CD38, anti-FOXP3 and anti-BCL6 antibodies and imaged them with confocal microscopy. CD38^bright^ cells, which likely belonged to the B lymphocyte lineage, were readily identifiable on stained sections, but so were CD38-expressing Tfr cells, which comprised 20% of the total Tfr pool (fig. S16, A and B). Regarding Tfr subset positions, 80% of CD38^+^Tfr cells localized within GCs, whereas most (85%) of CD38^-^ Tfr cells were identified outside GCs in a 50μm ring that contained some, but not most, of the follicular mantle (fig. S16, A and C). To determine Tfr subset locations within better-defined mantle, light, and dark zones, we analyzed CD38 expression on a CODEX-stained tonsil section. Unlike conventional confocal microscopy where Tfr cells were either clearly expressed CD38 or not, CD38 expression on CODEX-stained Tfr cells appeared more continuous (Fig. 6, C and D). To distinguish between CD38 positive and negative cells we set an expression threshold so that the proportion of CD38^+^ GC-resident Tfr cells identified by CODEX (59.5%; Fig. 6D) closely matched the proportion of counterparts definitively identified by confocal microscopy (58%; fig. S16D). Using this expression threshold, 52% of dark zone and 70.6% of light zone Tfr cells expressed CD38. In contrast, only 6.6% of mantle zone Tfr cells expressed CD38, a significantly different distribution (P<0.0001 by Chi-square tests for both comparisons; Fig. 6E). Hence, tonsillar CD38^+^Tfr cells preferentially reside in GCs nearby clonally related Tfh cells whereas CD38^-^Tfr cells reside mostly in the mantle, closer to Tregs.

To determine if CODEX-identified Tfr cells shared phenotypic features with CD25^hi^Tfh/Tfh or Treg lineage cells, we determined the proteins differentially expressed between CD38^+^ and CD38^-^Tfr cells (Fig. 6F). In agreement with flow cytometric analyses, proteins significantly upregulated in CD38^+^Tfr cells included Ki67, a prominent CD25^hi^Tfh cell marker, and several canonical Tfh cell molecules (BCL6, CXCR5, CD26 and SLAM) (*35*). In summary, tonsillar CD38^+^Tfr cells are clonally related to Tfh cells, retain a Tfh-like capacity for GC B-cell help, gain new regulatory functions through differentiation, and reside primarily within GC dark and light zones. In contrast, CD38^-^Tfr cells are clonally related to Tregs, appear highly specialized to suppress Tresp cell proliferation, and localize mostly to the follicular mantle.

## DISCUSSION

Herein, we describe previously unappreciated clonal, functional, and positional heterogeneity within the human tonsillar Tfr cell pool. Using *in-silico* transcriptomic trajectory projections, *in-vitro* tonsillar organoid lineage tracking, and *in-vivo TCRA/TCRB* sequencing we show human Tfh cells are not a terminally differentiated population. Rather, Tfh cells are capable of proliferating through a CD25^hi^BLIMP1^+^ intermediary stage before gaining FOXP3 expression and clear regulatory function as iTfr cells. Despite their considerable transformation, iTfr cells retain critical Tfh cell characteristics, like IL-21 secretion and GC-residence, that preserve the capacity for, and opportunity to provide, meaningful follicular B-cell help.

The existence of iTfr cells challenges early experiments in the Tfr field that demonstrated adoptively transferred murine Tregs, but not naïve T cells, entered the Tfr pool 7 to 11 days after vaccination (*7,12*). Our lineage tracking experiments using tonsillar organoids indicate that one week is likely too brief an interval for human naïve T cells to transition to Tfh cells, then to CD25^hi^Tfh intermediary cells and finally to Tfr cells. Indeed, more recently published longitudinal intra-vital imaging of mouse lymph nodes revealed FOXP3^+^ cells of Tfh origin accumulated in GCs 14 to18 days after vaccination and their influx heralded GC contracture (*11*). Our data suggest iTfr cells may arrive in the light zone at a similar GC life stage to suppress the same TCRαβ-matched Tfh clones they descended from. Since iTfr cells retain the ability to help B cells and permeate the dark zone, they may also maintain follicular output as GCs shrink. In contrast, nTfr cells are elite suppressors primarily positioned outside GCs but within follicular mantles suggesting a gatekeeping role that may include controlling autoreactive T cells at the T-B border.

Using non-biased, multi-omic analysis we identified CD38 as a practical cell surface marker to divide live, clonally divergent tonsillar Tfr subsets for downstream analyses, yet it is unclear if this molecule is important for iTfr cell differentiation or function. As CD38-deficient humans have not yet been identified, this line of inquiry may be best addressed using genetically manipulated animals. Although CD38 expression has been primarily described on human myeloid and B cells (*36*), recently a subset of GC-resident CD38 expressing CTLA4^hi^ Tfr cells were described within human mesenteric lymph nodes (*20*). Based upon our own microscopy findings, these cells and tonsillar iTfr cells may be analogous in both provenance and function.

While definitive identification of human Tfr cells requires intracellular FOXP3 staining, we did not clonally identify iTfr and nTfr cells nor sort-separate cells for functional analyses using differential FOXP3 expression since fixation/permeabilization would negatively affect mRNA integrity (*37*) and render cells inviable. Nevertheless, the clonal overlaps we identified between tonsillar Tfr and Treg cells and between Tfr and Tfh cells using cell surface stains were also reported in *Foxp3*^*gfp*^ reporter mouse lymph nodes (*12*). Instead of anchoring on FOXP3, our sorting strategy relied upon differential CD25, CXCR5, and PD1 expression to capture most Tfr cells without sacrificing purity. One described human Tfr subset definitively excluded by this approach was CD25^-^Tfr cells (*19*), which comprise <10% of the total tonsillar Tfr pool. Due to their omission, it is unclear if CD25^-^Tfr cells exist on the iTfr lineage continuum, the nTfr lineage continuum, or represent contributions from additional sources. One clue that suggests a relationship with Tregs is the reportedly high Helios expression by CD25^-^Tfr cells, a feature shared with nTfr cells. Future studies using more inclusive strategies will be required to definitively address CD25^-^Tfr cell provenance.

Finally, our study sought to describe the origins, functions and positions of Tfr cells in the secondary lymphoid tissues of healthy humans. Since lymph nodes and spleens are rarely excised from well subjects, we utilized tonsils from pediatric donors free of systemic inflammatory or immune deficiency diseases. If iTfr cells are generated in human lymph nodes, as they are in human tonsils and murine lymph nodes (*10,11*), iTfr/nTr imbalances may contribute to a spectrum of GC-relevant diseases including disorders caused by autoantibody production (i.e. systemic lupus erythematosus), poor vaccine responses (aging), or both (common variable immune deficiency; CVID). Indeed, increased circulating CD25^hi^Tfh cell frequencies and asymmetrically hyperplastic GCs have been uniformly described in CVID patients with co-morbid autoantibody-mediated cytopenias (*22,38*). Application of concepts, models and techniques described herein to diseased patient lymphoid tissues, rather than continued emphasis on studying their peripheral blood samples, may reveal unexpected pathophysiologies and provide new opportunities to intervene with precision therapies.

## MATERIALS AND METHODS

### Study design

The main aim of this study was to evaluate the contributions of Treg and Tfh cells to the human tonsillar Tfr pool. The study was conducted by multi-omic sequencing of 82,973 single CD4^+^ T cells from two immunocompetent tonsil donors. Microscopic imaging was performed on 10 tonsil tissue sections from six additional donors. Functional *in-vitro* analyses and independent confirmatory studies were conducted using material from 45 additional tonsil donors.

### Tonsillar T CD4^+^ cell subset preparation and sorting

Fresh tonsils were obtained as discarded surgical waste from de-identified immune-competent children undergoing tonsillectomy to address airway obstruction or recurrent tonsillitis. Tonsil donor mean age was 6 years and 53 % were male. Tissue collection was determined not to be human subjects research by the Children’s Hospital of Philadelphia Institutional Review Board. A single cell suspension of tonsillar mononuclear cells (MNCs) was created by mechanical disruption (tonsils were minced and pressed through a 70-micron cell screen) followed by Ficoll-Paque PLUS density gradient centrifugation (GE Healthcare Life Sciences). CD19-positive cells were removed (StemCell) and CD4^+^ T cells were enriched with magnetic beads (Biolegend) prior to sorting Tfh cells (CD4^+^CD25^-^ CXCR5^hi^PD1^hi^), CD25^hi^Tfh cells (CD4^+^CD25^hi^CXCR5^hi^PD1^hi^), Tfr cells (CD4^+^CD25^hi^CXCR5^+^PD1^int^), and Treg (CD4^+^CD25^hi^CXCR5^-^PD1^-^) on a BD FACSAria™ (BD Bioscience). Dead cells were excluded using LIVE/DEAD stain (Thermo Fisher Scientific). The gating strategy is shown in Fig.1A.

### CD4^+^ T cell subsets immunophenotyping

Tonsillar enriched CD4^+^ T cells were immunophenotyped using an LSRFortessa flow cytometer (BD Bioscience) with antibodies against AIOLOS (16D9C97), CD4 (OKT4), CD25 (BC96), CD38 (HIT2), CD39 (A1), CD127 (A019D5), CD130 (2E1B02s), CD161 (HP-3G10), CXCR5 (J252D4), FOXP3 (150D), GITR (108-17), HELIOS (22F6), IL-10 (JES3-9D7), Ki67 (Ki-67), LAG3 (11C3C65), and PD-1 (EH12.2H7) all from BioLegend; BCL6 (K112-91), CTLA4 (BNI3) and OX40 (ACT35) from BD Bioscience; BLIMP1 (IC36081A) from R&D systems; and IL-21 (3A3-N2), IL-17 (eBio64DEC17), and TIGIT (MBSA43) from Invitrogen. AIOLOS, BCL6, CTLA4, HELIOS, Ki67, LAG3, IL-10, IL-17, and IL-21 intracellular staining was performed after fixation and permeabilization with the Foxp3/Transcription Factor Staining Buffer Set (eBioscience) in accordance with the manufacturer’s instructions. To assess IL-10, IL-17, and IL-21 secretion by tonsillar CD4^+^, T cell subsets were rested overnight at 37°C then stimulated for 6 h with phorbol 12-myristate 13-acetate (25 ng/mL; Sigma) and ionomycin (1 mg/mL; Sigma) in the presence of Brefeldin A (5 μg/ml; BD Bioscience). Flow cytometric analyses were visualized with FlowJo software (TreeStar).

### CITE-Seq and ScRNA-Seq

Viable tonsillar Tfh, CD25^hi^Tfh, Tfr, and Treg cells were sorted from tonsils excised from two immune-competent male patients with obstructive sleep apnea. One patient was four and the other six-years-old. Transcriptome and TCR repertoire for each CD4^+^ T cell subset was obtained using a ScRNA-Seq approach. A CITE-Seq study was also developed on viable sorted Tfr cells from the 4-year-old donor. For the CITE-Seq sample preparation, Tfr cells were resuspended at 20 × 10^6^ cells/ml in CITE-seq staining buffer (BioLegend) and incubated with Human TruStain FcX Fc Blocking reagent for 10 min at 4°C to block nonspecific antibody binding. Following Fc blocking, cells were incubated with a pool of 139 antibodies conjugated to an antibody-derived tag (ADT), inclusive of seven isotype controls (TotalSeq-C Human Universal Cocktail, anti-human GITR (108-17) and anti-human CD130 (2E1B02); BioLegend) for 30 min at 4°C. After incubation, cells were washed three times with 3.5 ml of staining buffer to remove antibody excess. Cells were passed through a 40-μm filter to remove any cell clumps and resuspended in 10% FBS RPMI media at 1 × 10^6^ cells/ml for 10× Genomics 5’ single-cell RNA-seq. Next-generation sequencing libraries were prepared using the 10× Genomics Chromium Single Cell 5’ Library and Gel Bead kit v1 with Feature Barcoding Technology for Cell Surface Protein, per manufacturer’s “Chromium Single Cell V(D)J Reagent Kits” protocol. Libraries were uniquely indexed using the Chromium i7 Sample Index Kit, pooled, and sequenced on the Illumina NovaSeq 6000 sequencer (v1.5 chemistry) in a paired-end, single indexing run. Sequencing for each gene expression library targeted 20,000 mean reads per cell and each V(D)J library targeted 5,000 read pairs per cell. Data was then processed using the Cellranger multi pipeline (10x genomics, v5.0) for demultiplexing and alignment of sequencing reads to the GRCh38 transcriptome (10x Genomics v.3.0.0) and creation of feature-barcode matrices based on cell-associated cell barcodes across each paired gene-expression-VDJ library. CITE-Seq (ADT) data were identified for the Tfr library using the designated TotalSeq-C nucleotide sequences as an input to the cellranger multi pipeline. Data was aggregated using the cellranger aggr pipeline (10x Genomics, v.5.0). Secondary analysis was performed using the Seurat package version 4.1 (*39*) within the R computing environment. Gene barcode matrices were filtered to include cells expressing between 200-4000 genes and having mitochondrial content <20% of genes expressed. The filtered dataset was log-normalized, scaled, principal component analysis (PCA) performed, and clustered based on the top 2000 variable genes across the dataset. UMAP plots were used for visualization.

### V(D)J repertoire analysis

Immune profiling V(D)J data was added to the dataset by matching cell barcodes in the scRNAseq libraries. Shared stringent clonotypes between samples were defined as TCRα and TCRβ chains at the nucleotide level that were a direct match between at least two different T cell sub-populations in a minimum of two cells from each T cell population. scRepertoire within the R compute environment was used to evaluate clonal overlap, compute diversity metrics, and produce VDJ clonotype sharing visualizations between T cell subpopulations (*40*). Cell barcodes that had a TCRα, β chain that mapped to “NA” was excluded from downstream analysis. Further, the filterMulti command within the scRepertoire package was used to filter multi-mapping TCRα or TCRβ chains in each of the single cell VDJ libraries which identifies multimapping chains and provides a TCRα/β consensus call based on the highest expression for a single cellular barcode.

### Tonsillar organoid preparation

Once isolated, MNC were counted and resuspended in organoid media (RPMI with L-glutamine, 10% FBS, 2 mM glutamine, 1X penicillin-streptomycin, 1 mM sodium pyruvate, 1X MEM non-essential amino acids, 10 mM HEPES buffer, and 1 μg/ml of recombinant human B-cell activating factor [BioLegend]) at a concentration of 6 × 10^7^ cells per ml. As previously described by Wagar *et al*. 6 × 10^6^ MNC were transferred to permeable transwells (0.4-μm pore, 12-mm diameter; Millipore, (24)). Viable sorted Tfh and Treg cells from autologous tonsil donors were stained with carboxyfluorescein diacetate succinimidyl ester (CFSE; Thermo Fisher Scientific) or CellTrace Violet (CTV; Thermo Fisher Scientific), respectively, for tracking. 40 × 10^3^ Tfh cells or 10 × 10^3^ Tregs were added to sepreate organoids. Transwells were then inserted into standard 12-well polystyrene plates containing 1 ml of additional organoid media and placed in an incubator at 37 °C and 5% CO2. Organoid media was replaced every 3 days. On culture day 7, prior Tfh cells were identified according to the CFSE dye and evaluated for FOXP3, CTLA4, CD25, LAG3, and BLIMP1 expression. On culture day 8, prior Tregs were identified according to the CTV dye and evaluated for FOXP3 and CXCR5 expression.

### Treg cell suppression assays

CD4^+^CD45RO^-^CD25^-^ sorted naive responder T cells (5 × 10^3^) were labeled with CFSE and cocultured with an equal number of either Tfh, CD25^hi^Tfh, CD38^+^/CD38^-^Tfr, or Treg cells. Cultures were activated with anti-CD2/CD3/CD28 coated beads at a bead to cell of ratio of 1:1 (Miltenyi Biotec). Cocultures were stained for viability with the LIVE/DEAD Kit (Thermo Fisher Scientific), and the proliferation of viable responder T cells was determined by CFSE dilution at culture day 3.5 by flow cytometry.

### B-cell/T-cell cocultures and B-cell cultures

2.5 × 10^4^ viable CD19^+^CD21^+^CD38^+^IgD^-^ sorted GC B cells were cocultured with an equal number of viable sorted CD38^+^ or CD38^-^ Tfr cells. Cocultures were activated by addition of CD3/CD28–coated beads (Dynadeads, Sigma) at a ratio of 1 bead per T cell. Separatly, 5 × 10^4^ sorted viable GC B cells were cultured with megaCD40L at 1μg/ml (Enzo), recombinant IL-21 and/or recombinant both at 25ng/ml (R&D system). On day 7, culture supernatant IgG, and IgA concentrations were determined by ELISA.

### CODEX image processing and computational analysis

Raw TIF images were processed using MCMICRO (*41*), a Dockerized NextFlow pipeline implementing both BaSiC (Docker tag 1.0.1) for tile-level shading correction as well as ASHLAR (Docker tag 1.14.0) for inter-tile stitching and cross-cycle registration. The resulting OME.TIFF file output was segmented using a Dockerized implementation of Mesmer/DeepCell (Docker tag 0.10.0-gpu) (*42*) using the cycle 2 DAPI channel as “nuclear” input and a weighted sum of IgD, CD3, CD4, CD8, CD19, and CD95 channels as a composite “membrane” input. Briefly, each channel was re-scaled to the 99.9%-ile of pixel intensity, summed, and the result then rescaled to avoid saturation on a 16-bit color scale. Per-cell marker expression was then computed by the average raw pixel intensity of each marker over each cellular segmentation mask. The resulting cellular expression matrix was further processed and analyzed in R (v4.0.2) using Seurat (v4.0.4), with non-cellular artifacts manually flagged and removed based on cell size and autofluorescence outliers. Centered log ratio (CLR)-normalization was performed within each of 23 marker channels across cells prior to downstream principal component analysis (npcs=10), neighbor identification (dims=1:10), and cluster identification (resolution=0.4). Cellular clusters were named based on manual review of population-level marker distributions and spatial localization. Cellular neighborhood (CN) analysis was performed as previously described (*34*) using K=20 nearest neighbors. Due to the rarity of CD4^+^FOXP3^+^ populations in GCs, identification and annotation of such cells were manually curated by review of the raw imaging data. Prior to differential protein expression analysis, background subtraction was performed by subtracting the cycle 1 “blank” signals for the respective imaging channel from each marker. CD38 expression patterns were independently confirmed by high-resolution confocal imaging to ensure CD38 signals identified by CODEX imaging were not highly influenced by membrane spillover (43) from B cells onto T cells.

### Immunostainings and fluorescence microscopy

Tonsils were frozen in Tissue-Tek O.C.T™ Compound (Sakura Finetek) and 6μm-section were prepared and mounted on SuperFrost™ Plus glass slides (Fisher Scientific), fixed in cold acetone for 20 min and air dried. Rehydration and washing in Tris Buffered Saline (TBS) followed. Non-specific antibody binding was avoided using Blocking Reagent™ (Perkin Elmer) for 30 min. Sections were then incubated overnight with anti-human CD38 (EPR4106, abcam), FOXP3-eFluor570 (236A/E7, Invitrogen) BCL6-AF647 (K112-91, BD Bioscience) antibodies at 4 °C overnight. To detected CD38 protein, sections were incubated with anti-rabbit AF488 (Invitrogen) antibodies for 2h at room temperature. After fixation with paraformaldehyde 4% (Electron Microscopy Sciences) for 20 min, slides were then stained with DAPI (1 μg/ml, Sigma) for 15 min at room temperature and mounted with fluorescent mounting medium (DAKO). All images were acquired using a Leica TCS SP8 Confocal and analyzed using ImageJ software.

### Statistical analyses

Prism v9 (GraphPad) was used to perform non-parametric Mann-Whitney tests for analysis of immunophenotypic data, functional assays. Chi-square and Spearman correlation R coefficient tests were performed for Tfr cell localization analysis. Non-parametric statistical correlation analysis was performed as previously described (https://github.com/wherrylab/statistics_code) for differential protein expression analysis of CODEX-stained tonsillar tissue. Differential expression analysis was performed in the scRNAseq and CITE-seq data using a Wilcoxon Rank Sum test and a Bonferroni correction to account for multiple comparisons.

## Acknowledgments

We thank the subjects and their families. We thank the Children’s Hospital of Philadelphia’s Division of Otolaryngology, Center for Applied Genomics and Flow Cytometry Core for pediatric tonsil samples, next-generation sequencing and technical support, respectively. We thank the Sayer family for financial support.

## Funding

This work was supported by grants from the National Institutes of Health,

National Institute of Allergy and Infectious Diseases AI146026 (NR)

National Institute of Allergy and Infectious Diseases AI155577 (EJW)

National Institute of Allergy and Infectious Diseases AI115712 (EJW)

National Institute of Allergy and Infectious Diseases AI117950 (EJW)

National Institute of Allergy and Infectious Diseases AI108545 (EJW)

National Institute of Allergy and Infectious Diseases AI082630 (EJW)

National Institute of Allergy and Infectious Diseases CA210944 (EJW)

National Institute of Allergy and Infectious Diseases AI114852 (RSH)

National Heart, Lung, and Blood Institute R38 HL143613 (DAO)

National Heart, Lung, and Blood Institute T32 HL 7775-28 (JPB)

National Cancer Institute T32 CA009140 (DAO)

Additional funding from

Parker Institute for Cancer Immunotherapy (EJW)

Parker Institute for Cancer Immunotherapy, Parker Bridge Fellow Award (DAO)

Gray Foundation (DP)

Chan Zuckerberg Initiative Pediatric Networks for the Human Cell Atlas (NR)

## Author contributions

Conceptualization: CLC, DAO, NR

Methodology: CLC, DAO, RSH, JPB, DP, MG, NR

Formal analysis: CLC, NDL, DAO, JG, MG

Investigation: CLC, DAO, RSH, NDL, ECC, JPB, DP, LVS, AVCK, SY

Resources: KBZ, SDH, HH, EJW

Writing – original draft: CLC, NR

Writing – review & editing: all authors

Visualization: CLC, DAO, NDL, AVCK, MG

Supervision: HH, EJW, NR

Project administration: NR

Funding acquisition: DAO, RSH, JPB, DP, EJW, NR

## Data and materials availability

All data are available in the main text or the supplementary materials.

## Supplementary materials

**Supplementary Figure 1.**
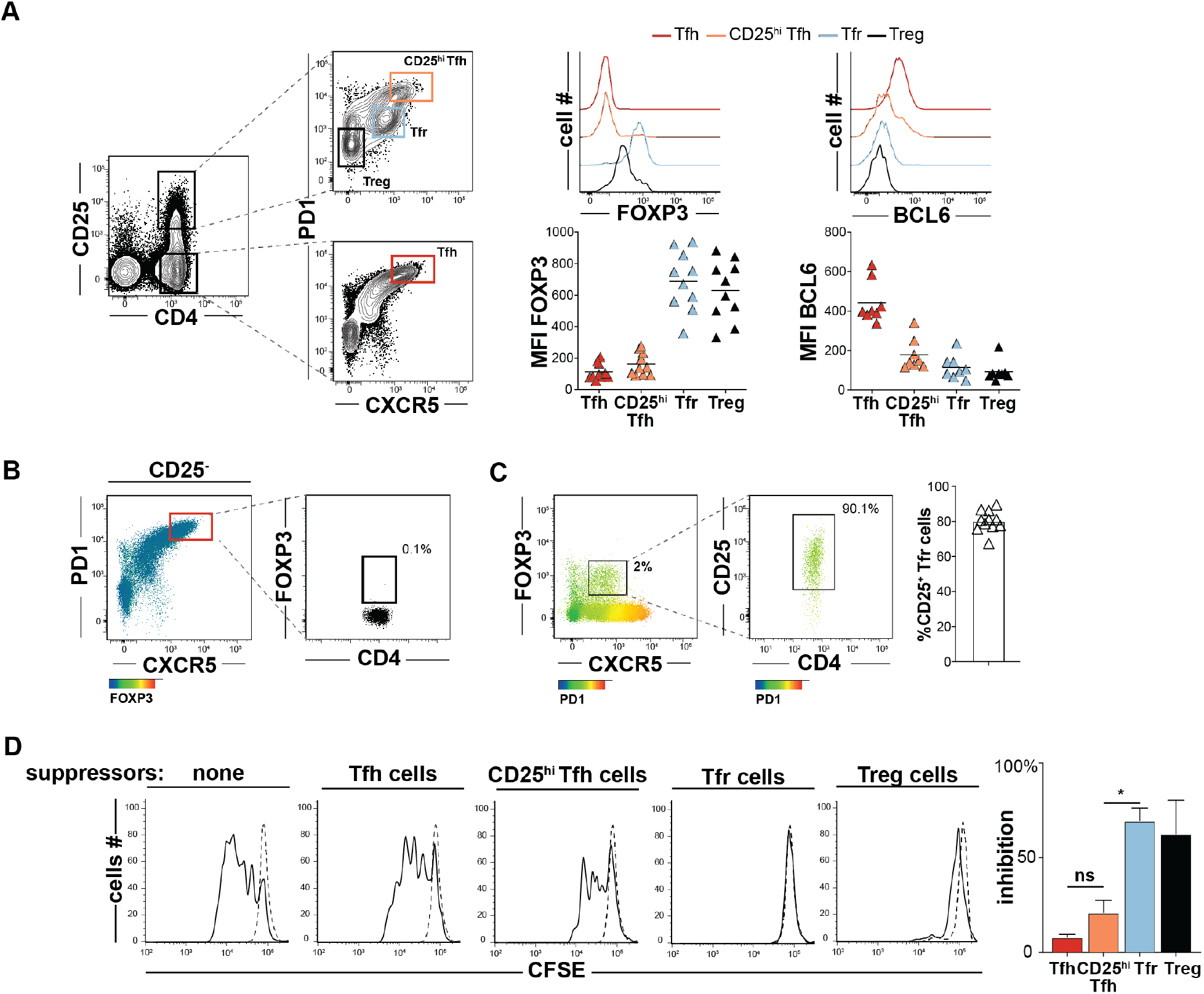
Treg and follicular T helper cell sorting strategy. (**A**, left) Treg (CD4^+^CD25^hi^CXCR5^-^PD1^-^, black), Tfr (CD4^+^CD25^hi^CXCR5^+^PD1^int^, light blue), CD25^hi^Tfh (CD4^+^CD25^hi^CXCR5^hi^PD1^hi^, orange) and Tfh (CD4^+^CD25^-^CXCR5^hi^PD1^hi^, red) gates are displayed on T cells from a representative tonsil donor. (**A**, right) Histograms and plots display FOXP3 and BCL6 mean fluorescence intensities (MFIs) on T helper subsets from a representative and all tonsil donors (n=6-8), respectively. Two-dimensional dot plots show (**B**) FOXP3 expression on cells in the Tfh gate and (**C**) PD1 expression on FOXP3^+^CXCR5^+^CD25^hi^ Tfr cells from a representative donor and all donors (n=11). (**D**, left) Representative histograms of CFSE-labeled T-cell responders (Tresp, CD4^+^CD25^-^CD45RO^-^) stimulated (solid line) or not (dashed line) in coculture with indicated tonsillar T helper cell subsets are displayed. Bar graph (right) represents mean percent inhibition relative to unstimulated Tresp cells from three tonsil donors. Error bars means ± SEMs. *, P<0.05 by Mann-Whitney U test.

**Supplementary Figure 2.**
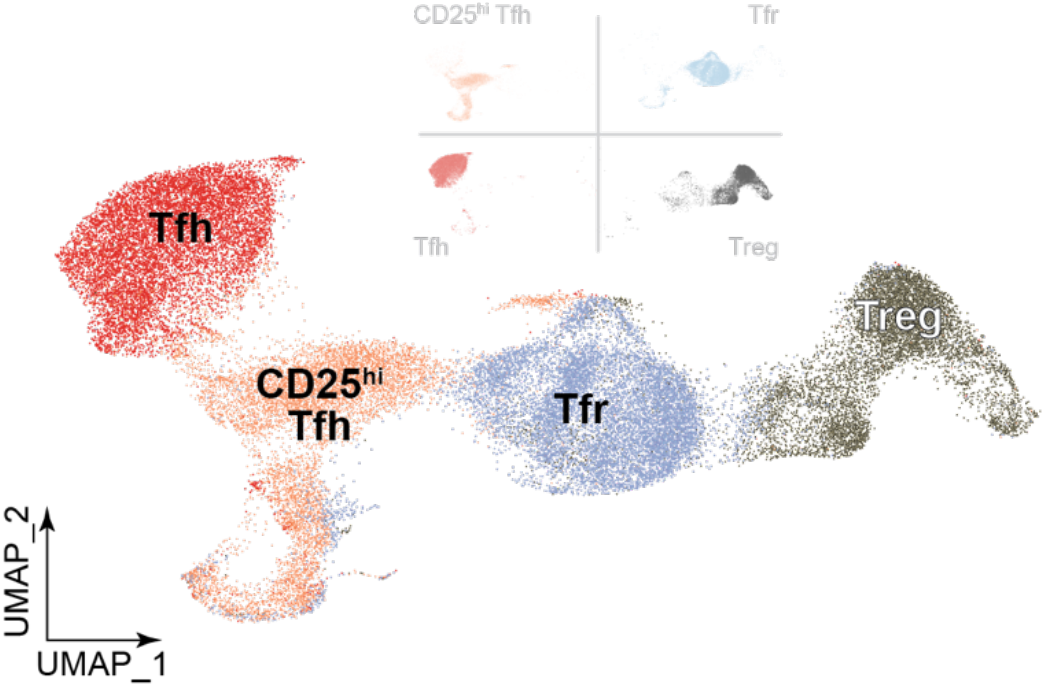
Dimensionally reduced transcriptomes of indicated TC341 cell subsets are displayed as a UMAP.

**Supplementary Figure 3.**
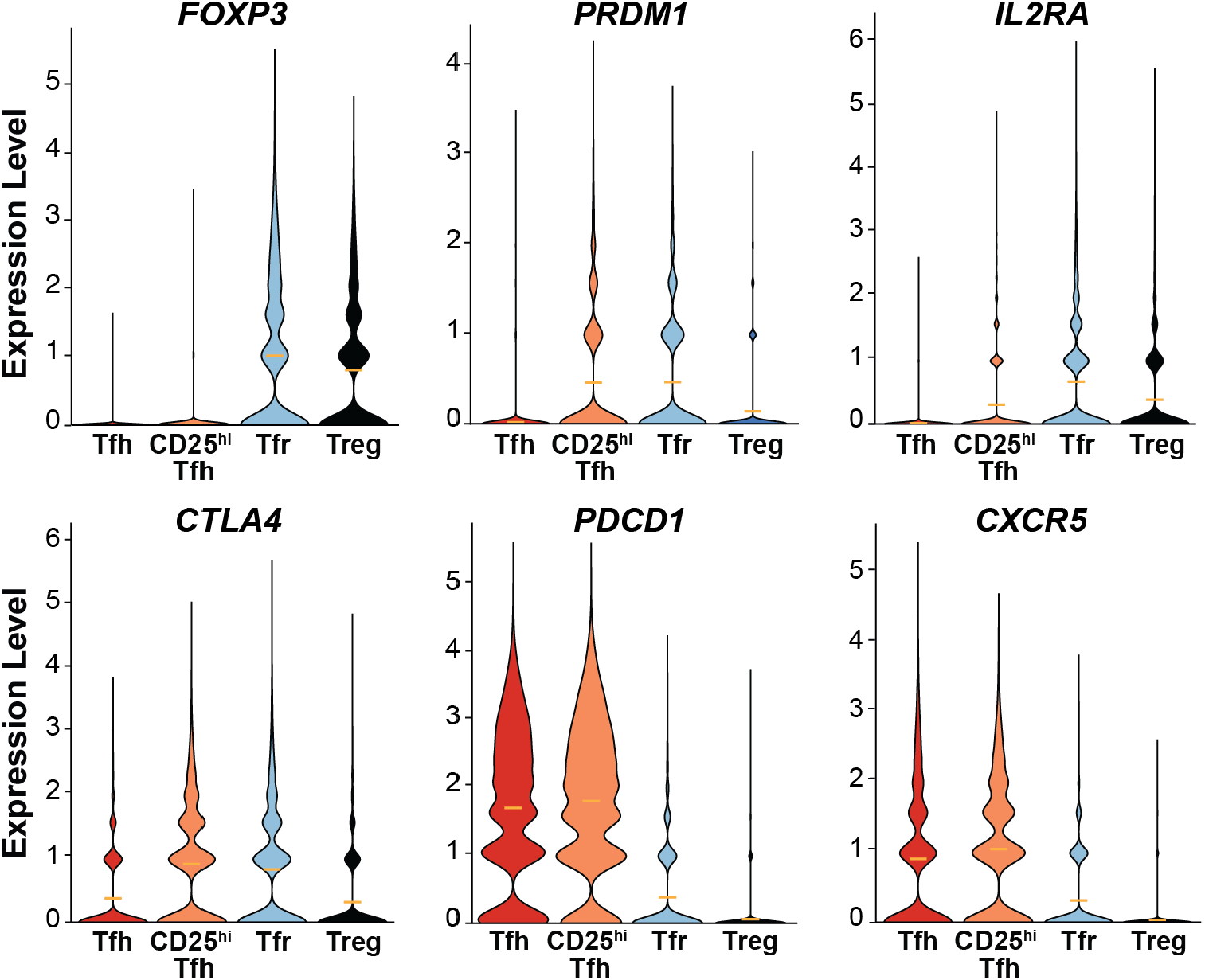
Violin plots display differentially expressed transcripts from pooled tonsil donor cells (TC174 and TC341).

**Supplementary Figure 4.**
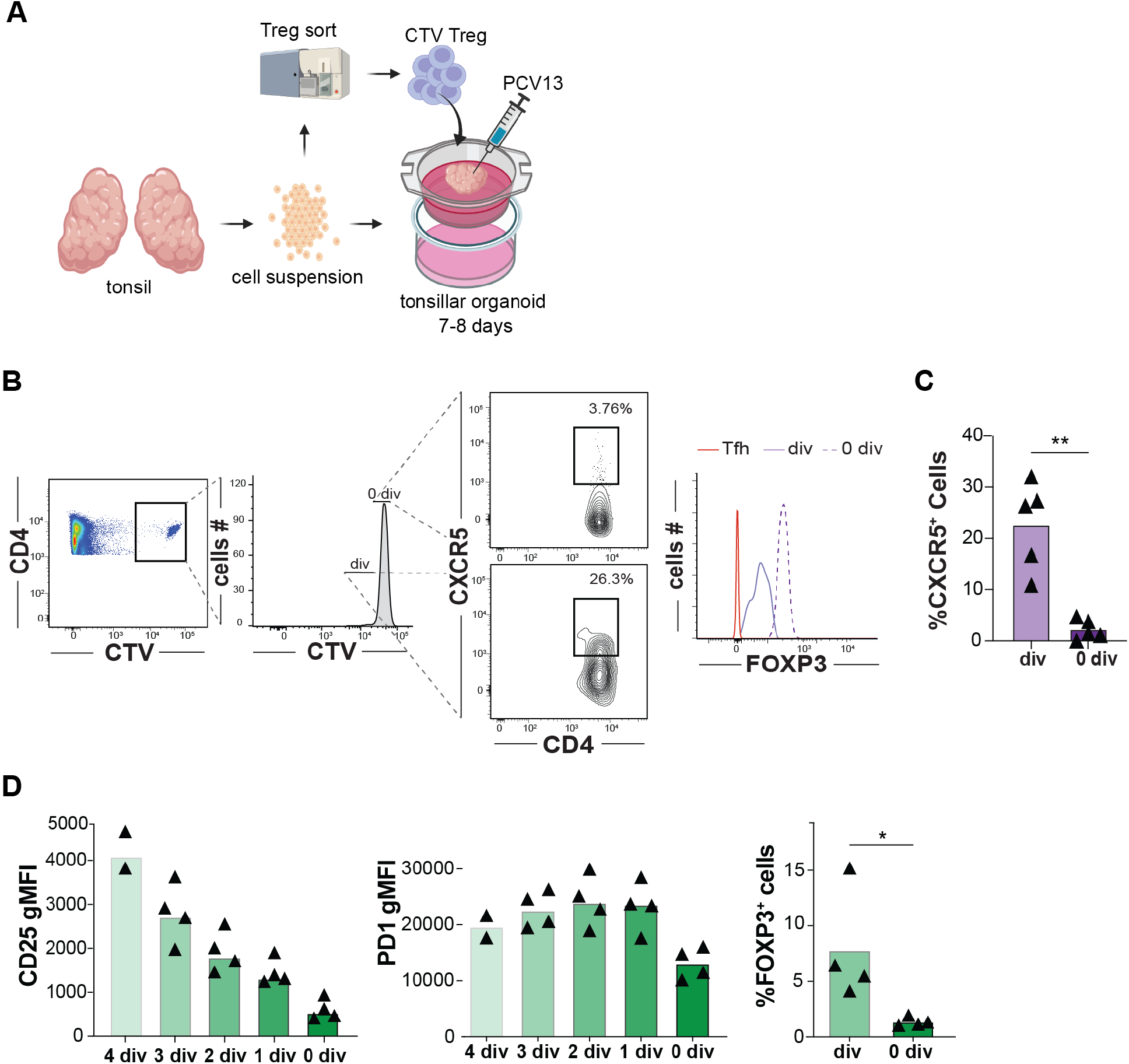
A strategy to track Tregs within vaccinated tonsillar organoids and measure differential protein expression on organoid incorporated Treg and Tfh cells. (**A**) is depicted by cartoon. (**B**) Divided (div) Cell Trace Violet™ (CTV) stained Tregs from a representative organoid experiment differentially upregulate CXCR5 at day 7-8 compared with undivided (0 div) counterparts yet maintain FOXP3 expression. (**C**) CXCR5 expression of divided and undivided cells from five independent experiments are shown. (**D**) CD25, PD1, and FOXP3 expression by divided and non-divided CFSE stained-Tfh cells are displayed 7 days after PCV13 vaccination in tonsil organoids (n=4) *, *P*<0.05 ; **, *P*<0.01 by Mann-Whitney U test.

**Supplementary Figure 5.**
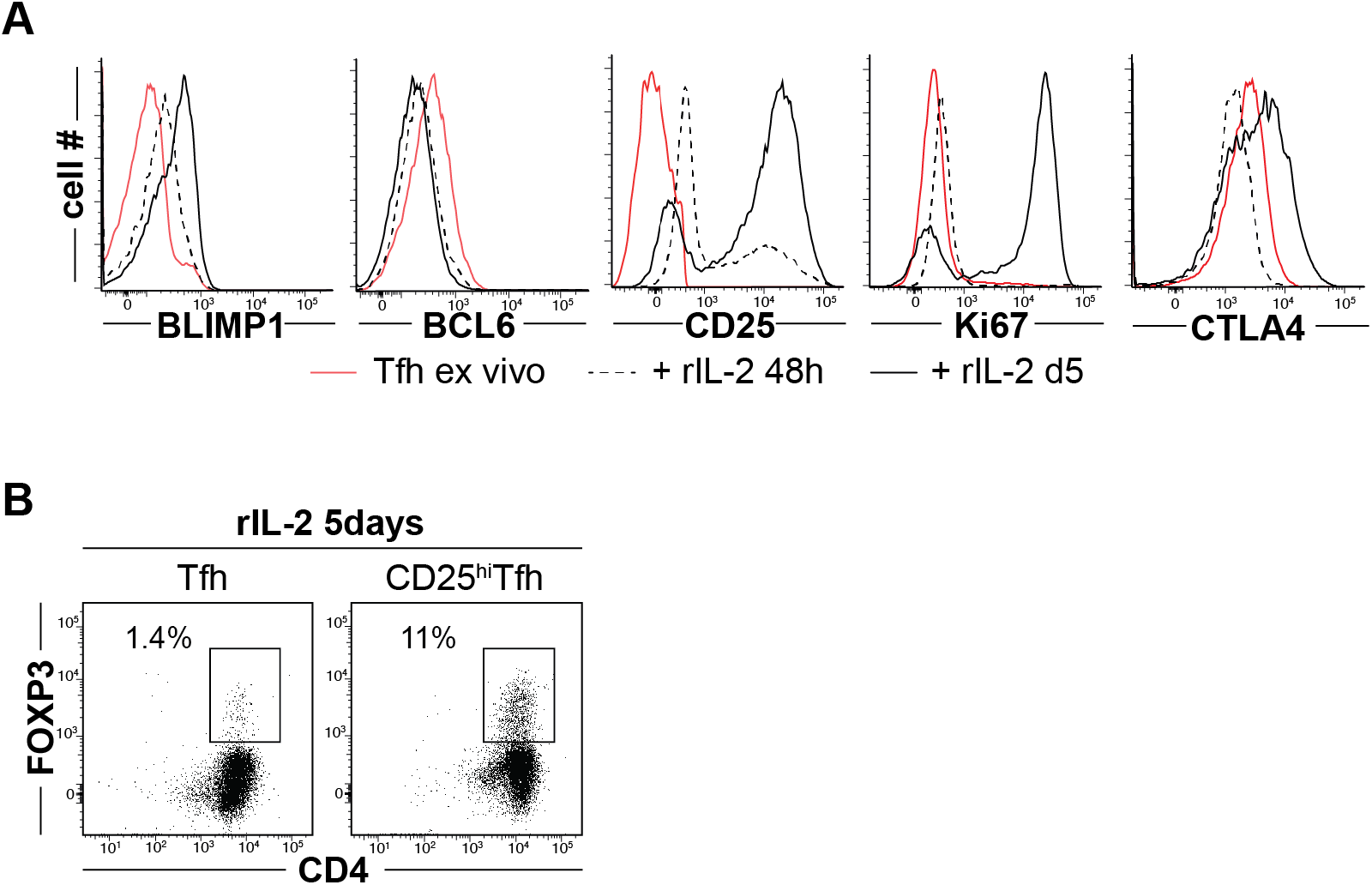
In-vitro Tfh and CD25hiTfh cell differentiation experiments. (**A**) Representative histograms of BLIMP1, BCL6, CD25, Ki67 and CTLA4 expression by Tfh cells before (red), after 48hours (dashed line) and five days (solid line) of recombinant IL2 treatment. (**B**) Representative dot plot of Tfh (left) and CD25^hi^Tfh cell (right) FOXP3 expression after five days of recombinant IL2 (rIL-2) treatment.

**Supplementary Figure 6.**
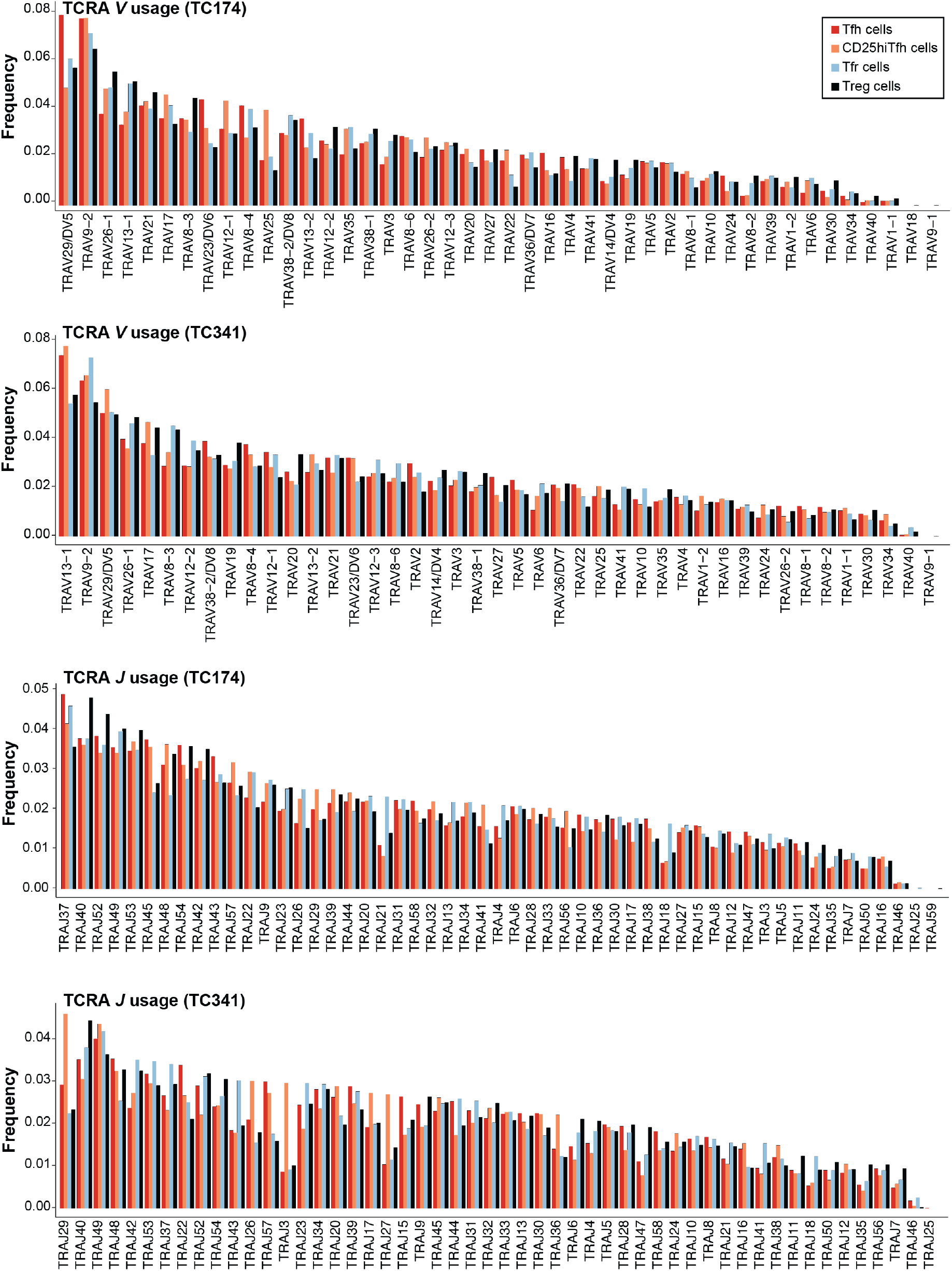
TCRA gene segment usage of TC174 and TC341. Tfh cells (red), CD25^hi^Tfh cells (orange), Tfr cells (light blue) and Tregs (black) TCR alpha chain *V* and *J* usage frequencies in tonsil donors TC174 and TC341.

**Supplementary Figure 7.**
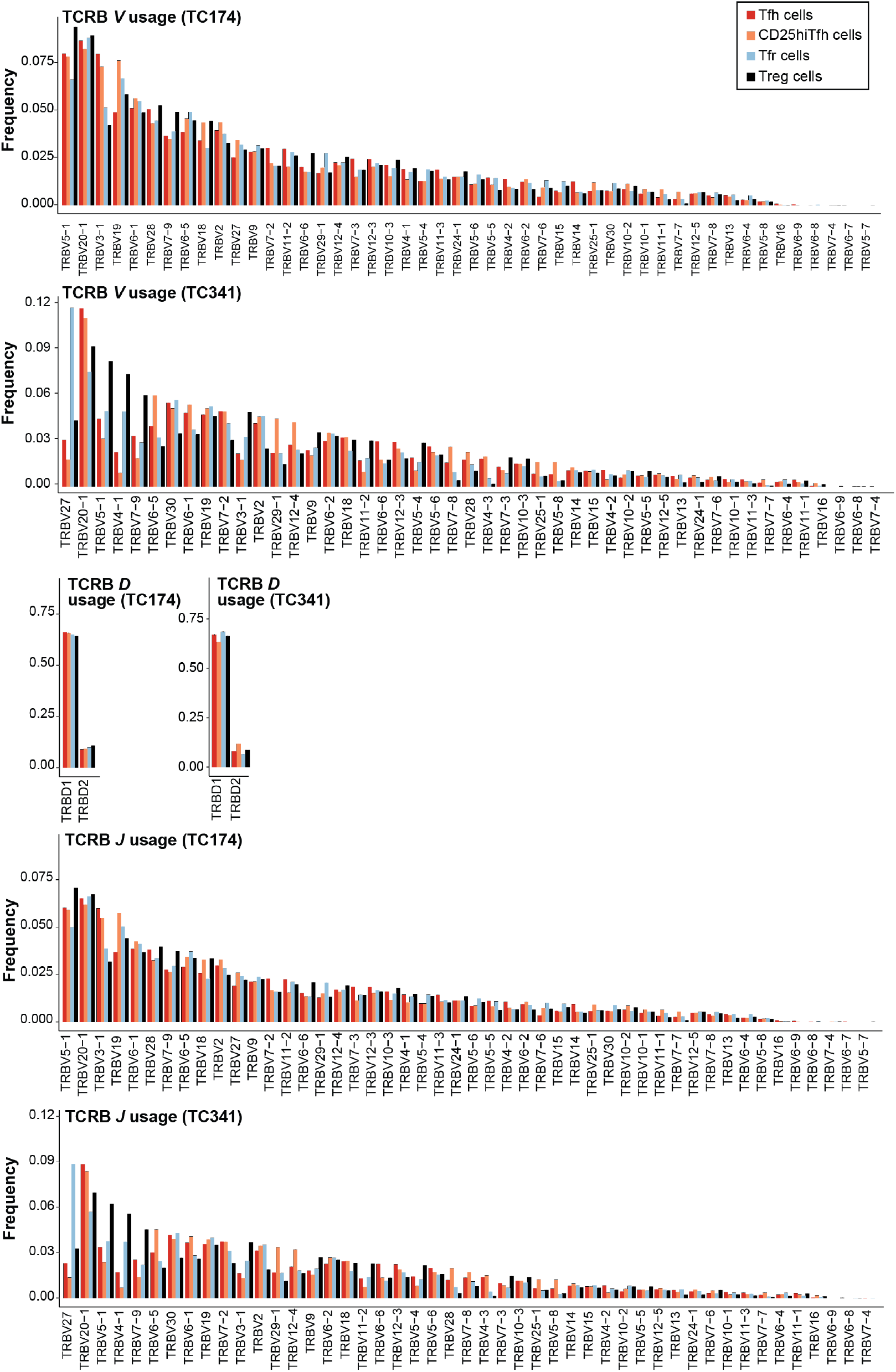
TCRB gene segment usage of TC174 and TC341. Tfh cells (red), CD25^hi^Tfh cells (orange), Tfr cells (light blue) and Tregs (black) TCR beta chain *V, D* and *J* gene segment usage frequencies in tonsil donors TC174 and TC341.

**Supplementary Figure 8.**
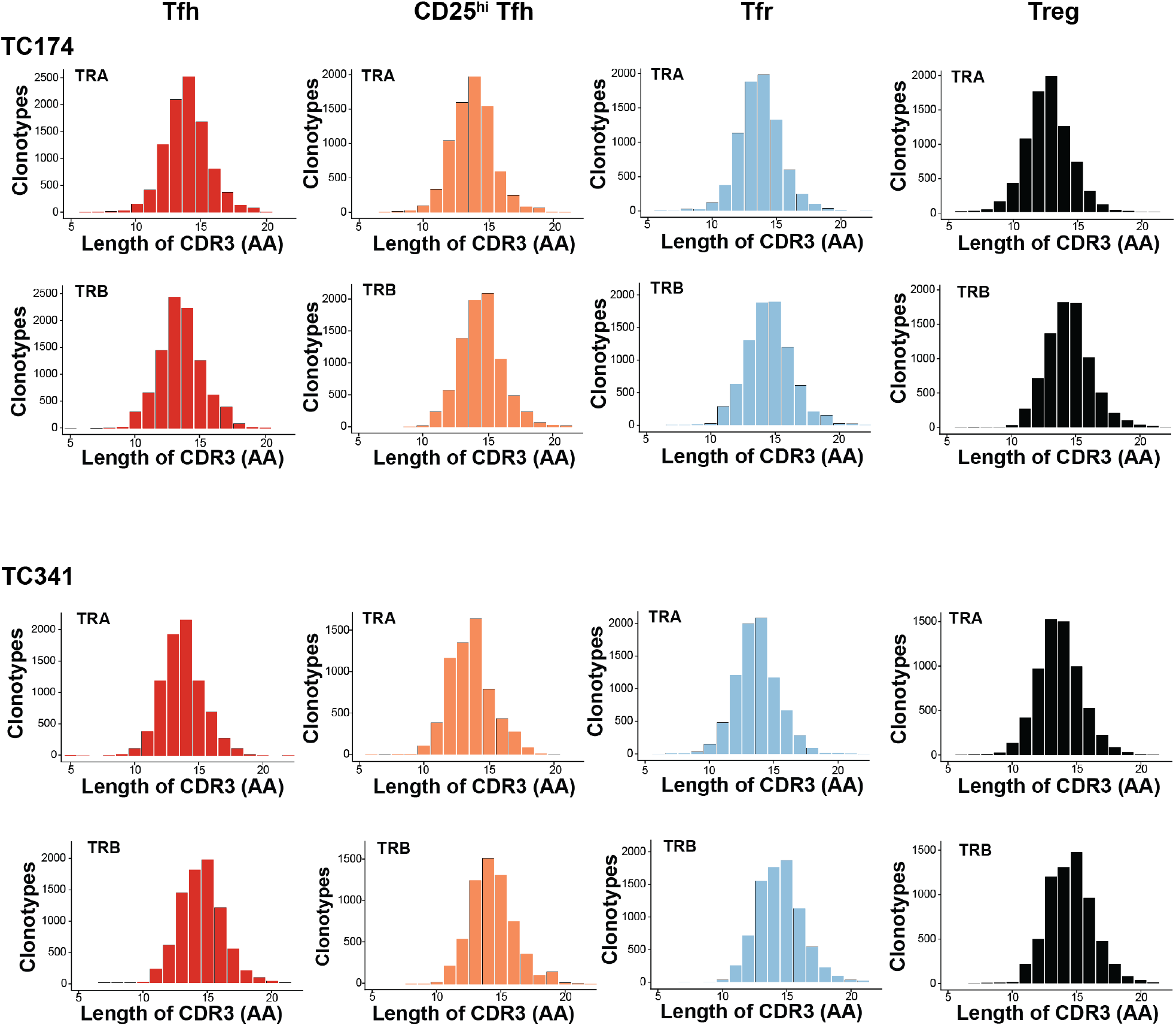
TCR CDR3 length distributions of TC174 and TC341. Distribution of TCRA (above) and TCRB (below) CDR3 segments of indicated T-cell subsets (Tfh, red; CD25^hi^Tfh, orange; Tfr, light blue and Treg, black) from two TC174 and TC341.

**Supplementary Figure 9.**
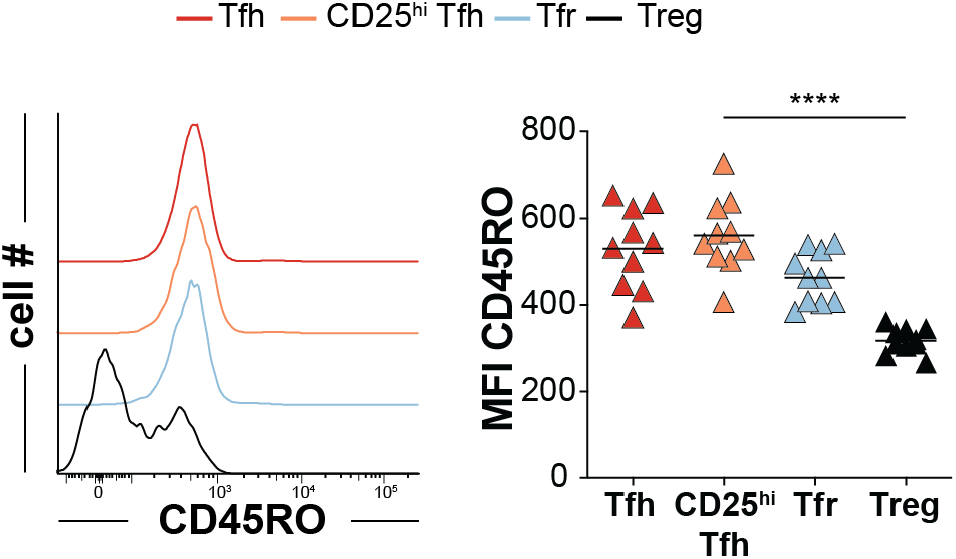
Differential CD45RO expression by CD4+ T cell subsets. Tfh cells (red), CD25^hi^Tfh cells (orange), Tfr cells (light blue) and Tregs (black) CD45RO are displayed as (left) histograms from a representative tonsil donor (right) and all tonsil donors (n=10). ****, *P*<0.0001 by Mann-Whitney U test.

**Supplementary Figure 10.**
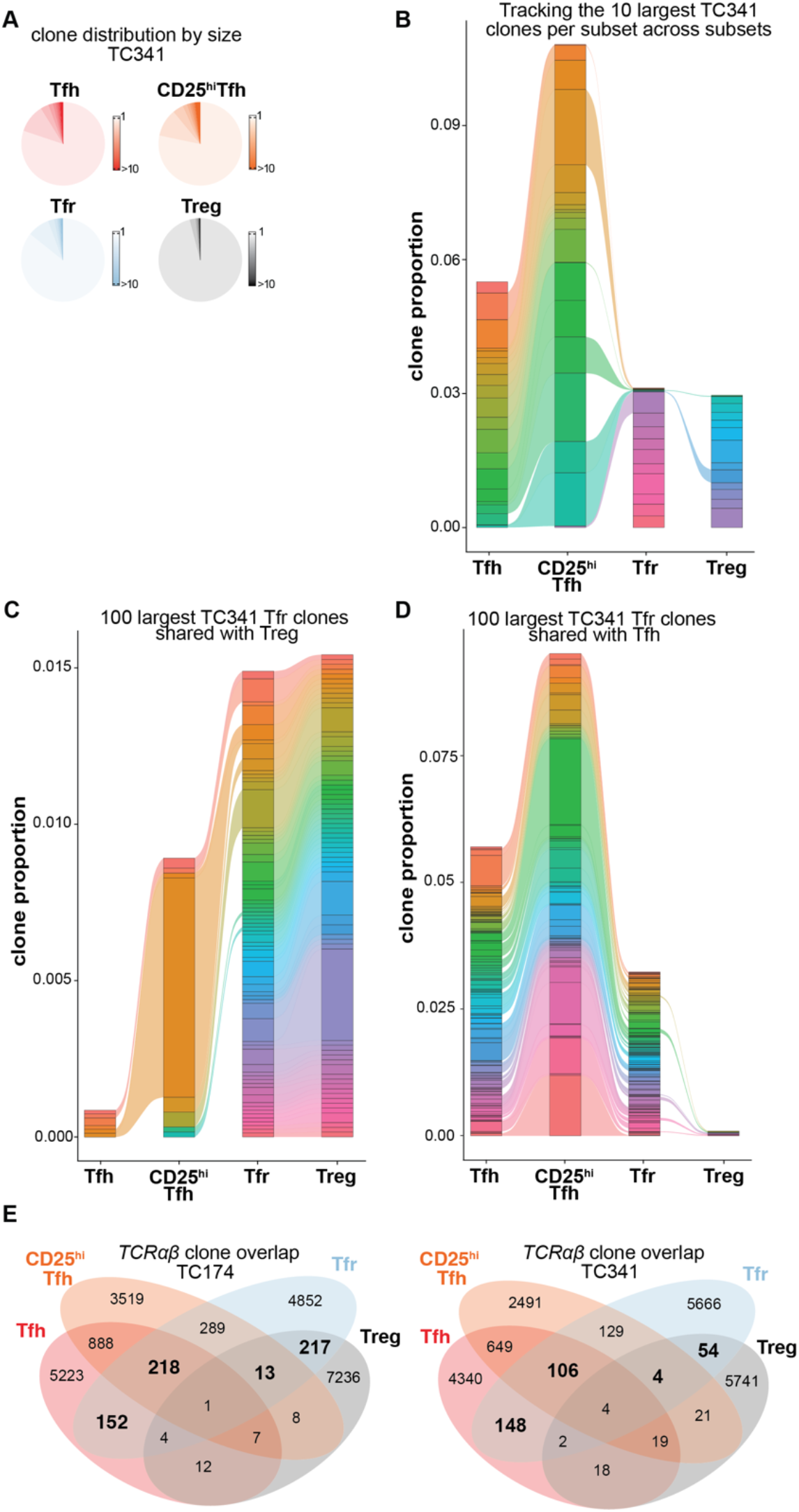
Clone size distribution and clone sharing between T helper subsets from TC174 and TC341. (**A**) Pie charts indicate subset-specific clone size distributions (cells per clone) for TC341. (**B**) Tracking the top ten largest clones per subset across other subsets. (**C**) Tracking the top 100 largest clones shared between Tfr and Tfh cells and (**D**) between Tfr and Tregs across other subsets. More than ten clones or 100 clones per subset are sometimes displayed to account for ranking ties and clone sharing across subsets. **E**) Venn diagrams depict clone sharing between T helper cell subsets from TC174 or TC341.

**Supplementary Figure 11.**
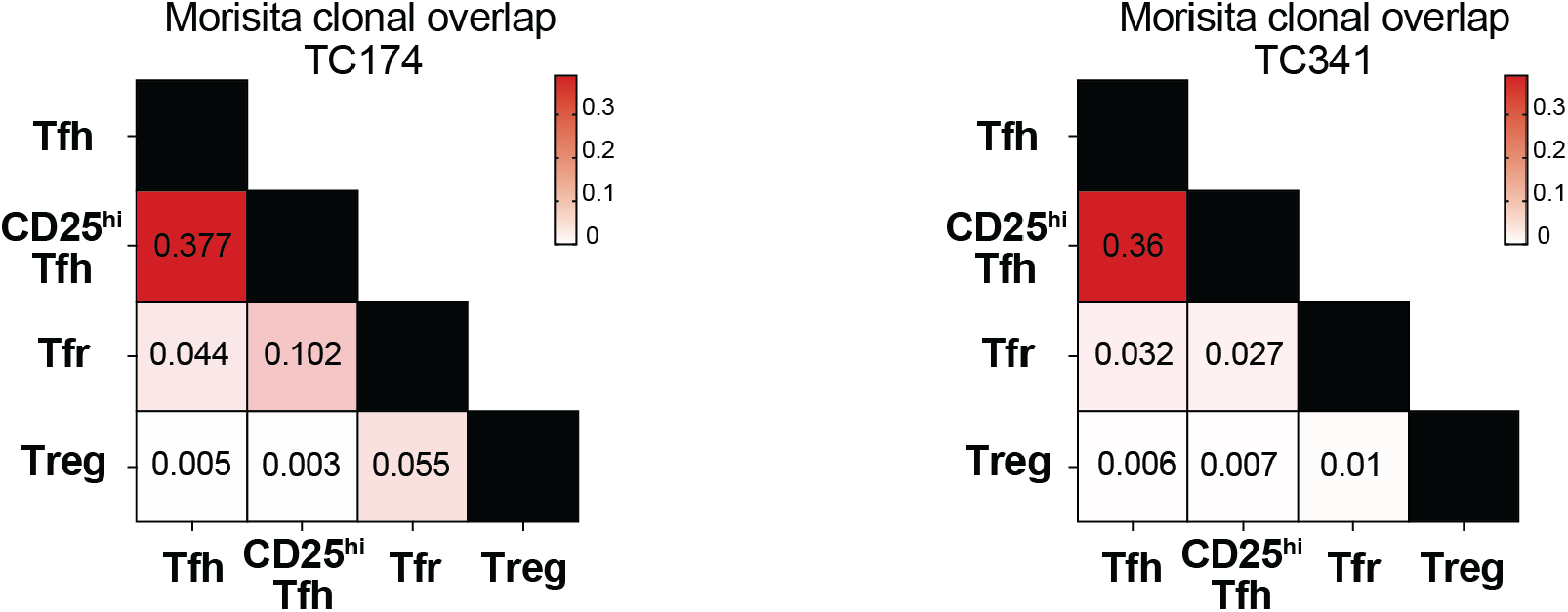
Morisita clonal overlap frequencies of TC174 and TC341. T helper subsets from TC174 (left) and TC341 (right). Red shading indicates more clonal sharing.

**Supplementary Figure 12.**
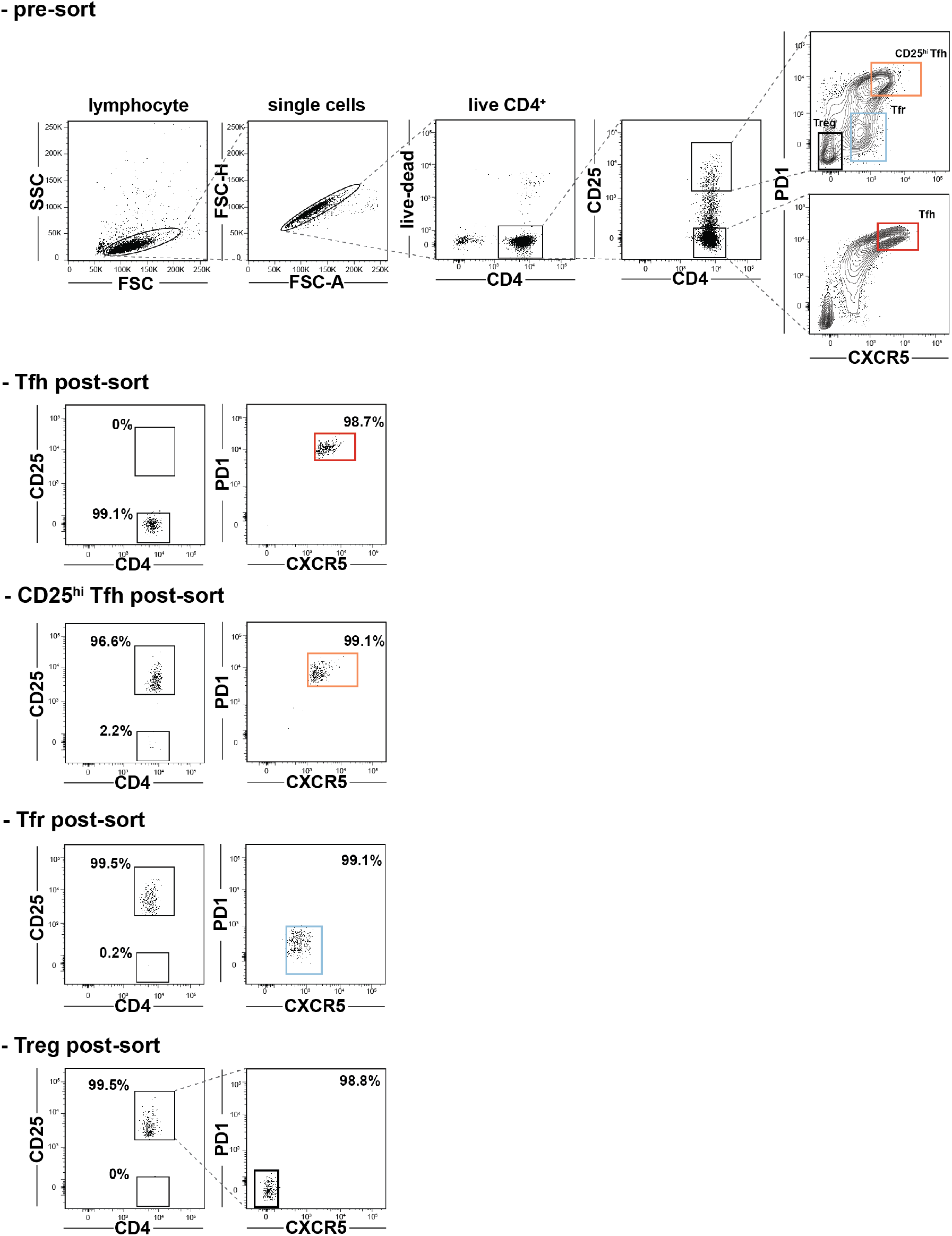
Pre and post-sort cell purity assessments. (above) Sorting strategy to sort Tfh, CD25^hi^Tfh, Tfr, and Treg cells. (below) Post-sort purities of the same subsets from a representative tonsil donor are displayed.

**Supplementary Figure 13.**
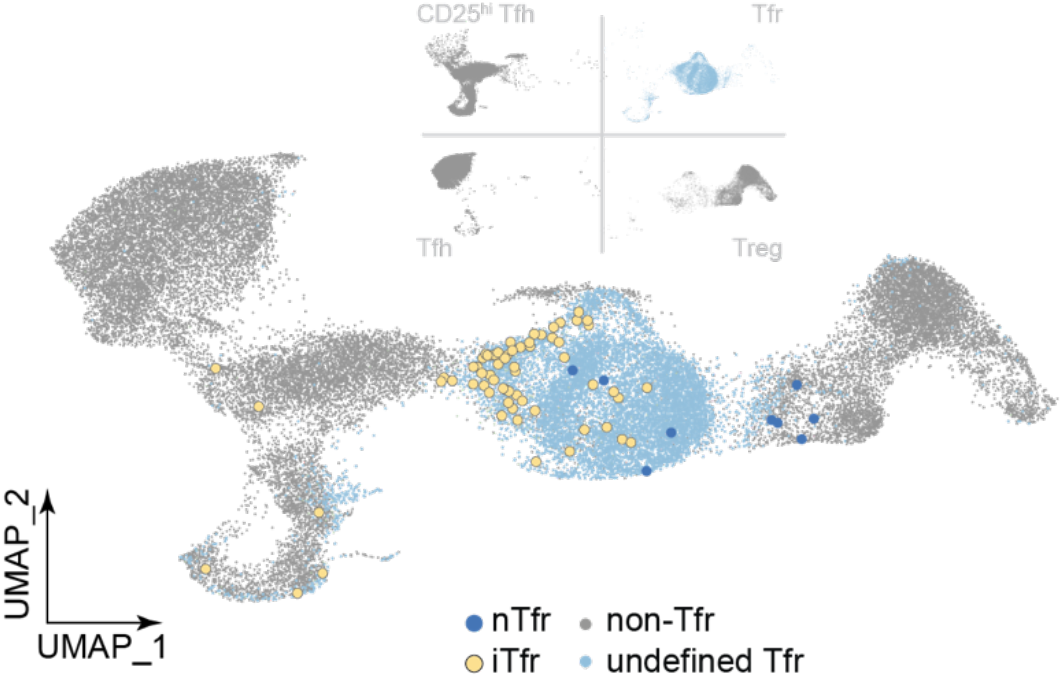
iTfr and nTfr localizations on the TC341 UMAP. iTfr (yellow) and nTr (dark blue) cell positions are indicated relative to clonally-undefined Tfr cells (light blue) and non-Tfr cells (gray).

**Supplementary Figure 14.**
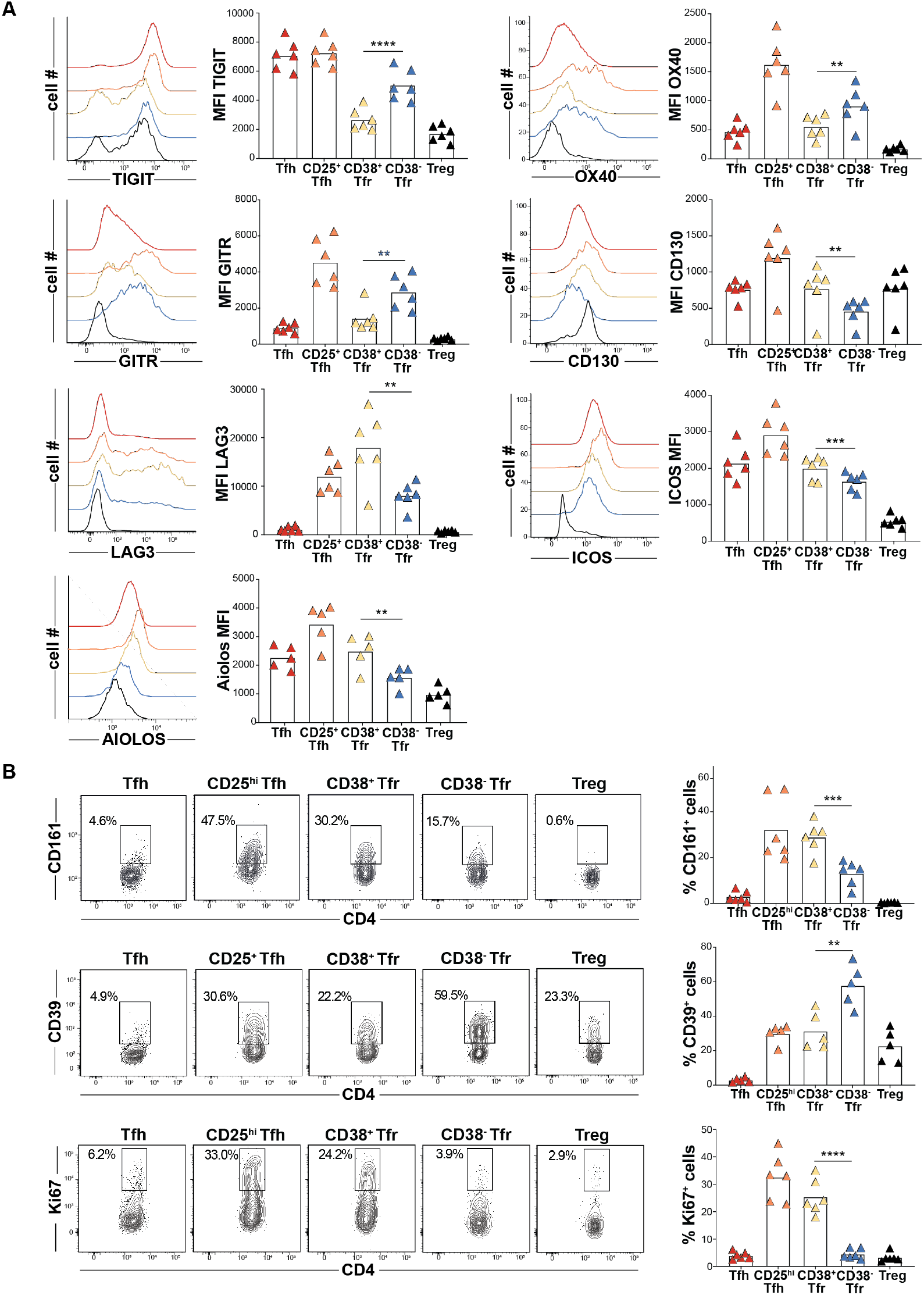
CD38^+^ and CD38^-^ Tfr cell extended immunophenotypes. (**A**) Histograms display TIGIT, OX40, GITR, CD130, LAG3 ICOS and AIOLOS expression by indicated T helper cell subsets from a representative tonsil donor (left) and bar graphs shows geometric mean fluorescence intensities (gMFIs) from counterpart cells fom five to six tonsil donors. (**B**) Dot plots present CD161, Ki67 and CD39 frequencies of indicated subsets from a representative tonsil donor (left) and five to six tonsil donors (right) **, *P*<0.01; ***, *P*<0.001; ****, *P*<0.0001 by Mann-Whitney U tests.

**Supplementary Figure 15.**
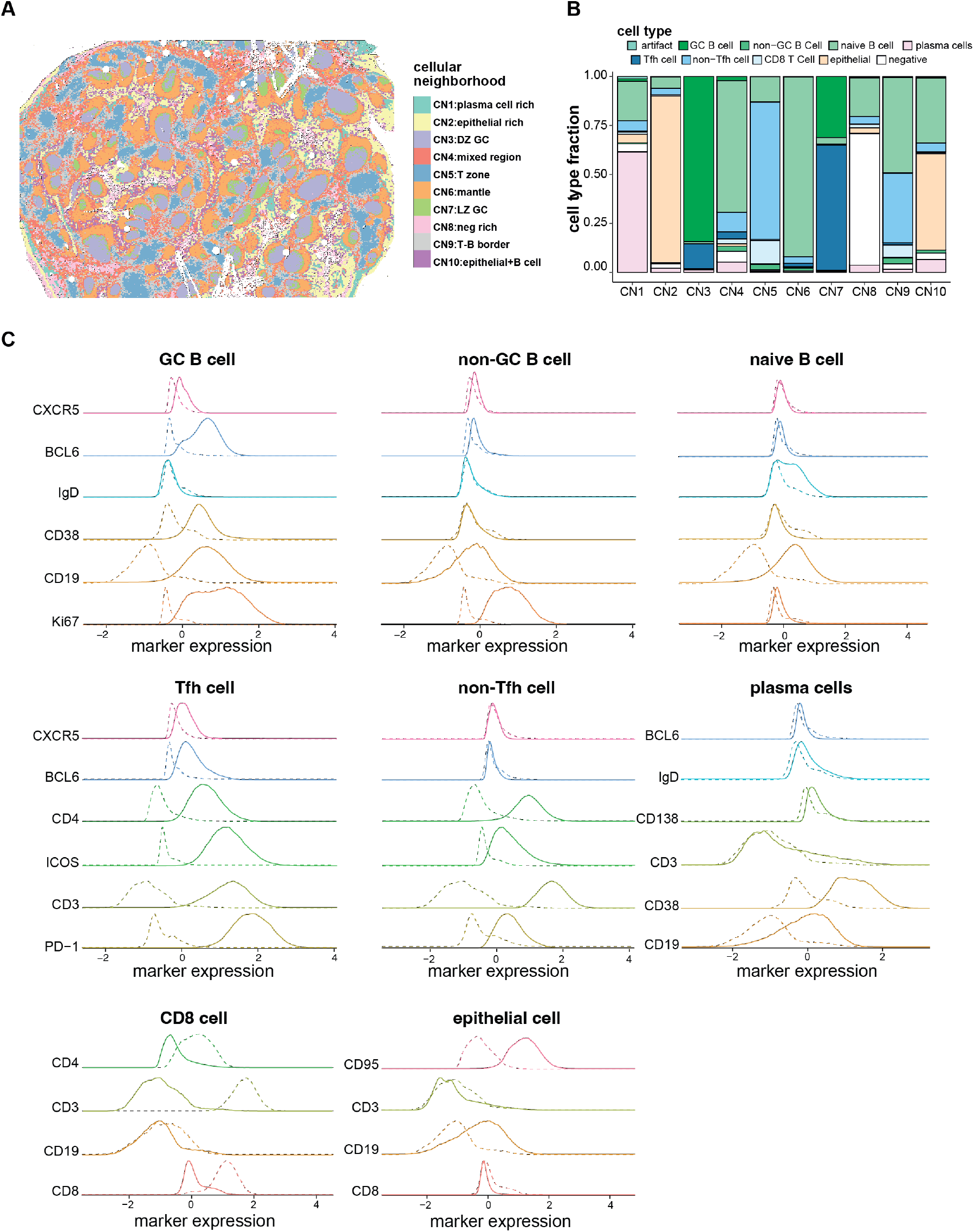
Tonsil cellular neighborhoods (CNs) and cell type profiles. (**A**) CN positions, (**B**) cell type compositions by CN, and (**C**) cell type protein expression profiles of a CODEX-stained tonsil section are displayed.

**Supplementary Figure 16.**
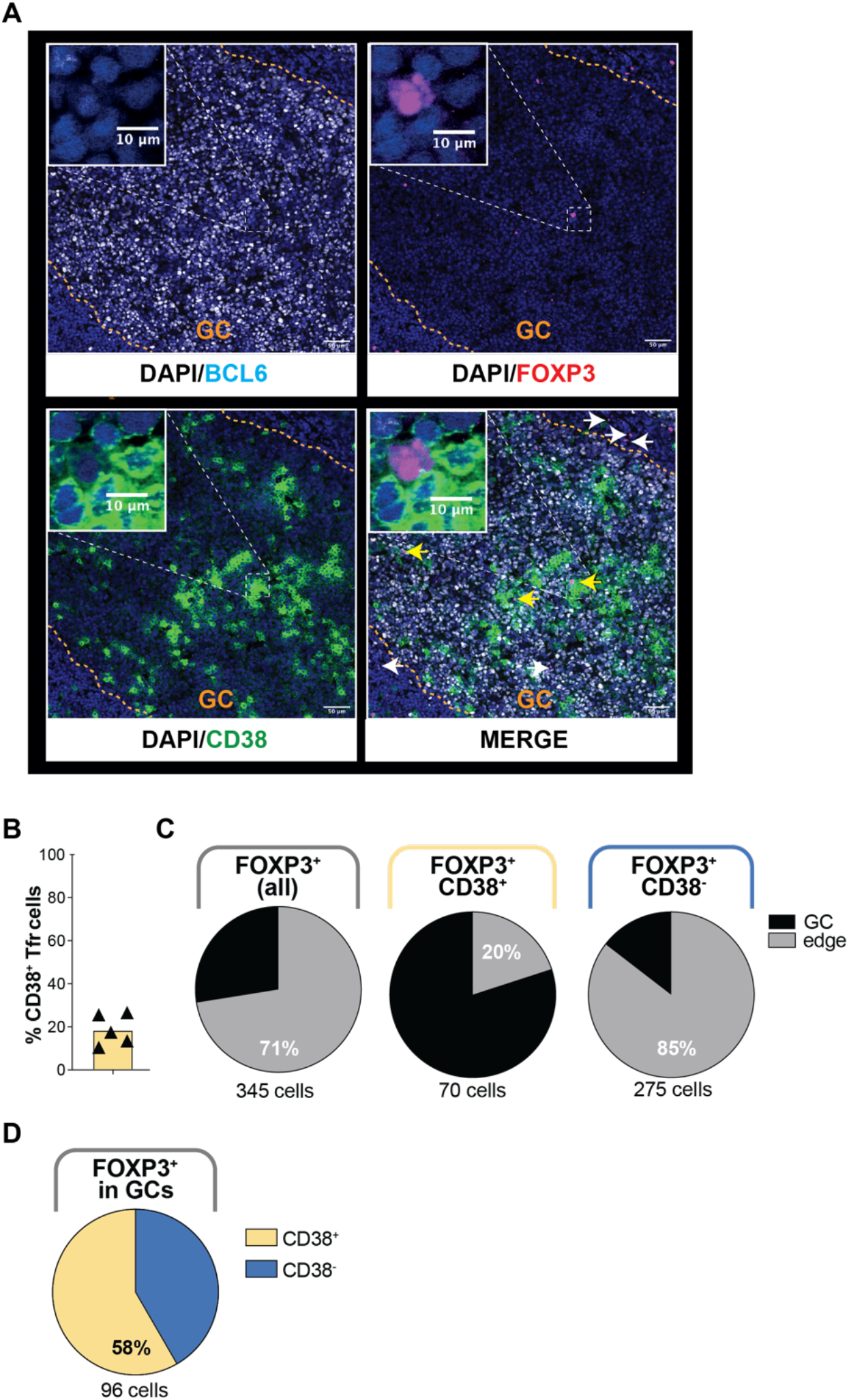
Confocal microscopy of stained tonsillar germinal centers (GCs). (**A**) BCL6, FOXP3, CD38 and DAPI stains of a representative tonsil section are displayed separately and merged. The positions of CD38^+^FOXP3^+^ cells (yellow arrows) and CD38^-^FOXP3^+^ cells (white arrows) are indicated. White dashed squares contain one representative CD38^+^FOXP3^+^ cell magnified in insets at the top left corner. (**B**) Frequencies of GC-associated FOXP3^+^ cells that express CD38, or not, from five tonsil donors are shown. (**C**) Proportions of FOXP3^+^, CD38^-^FOXP3^+^, and CD38^+^FOXP3^+^ cells located within BCL6-defined GCs and outside them in the surrounding 50μm GC edge region and (**D**) Proportions of CD38^-^FOXP3^+^ and CD38^+^FOXP3^+^ cells located within BCL6-defined GCs are displayed.

**Supplementary Figure 17.**
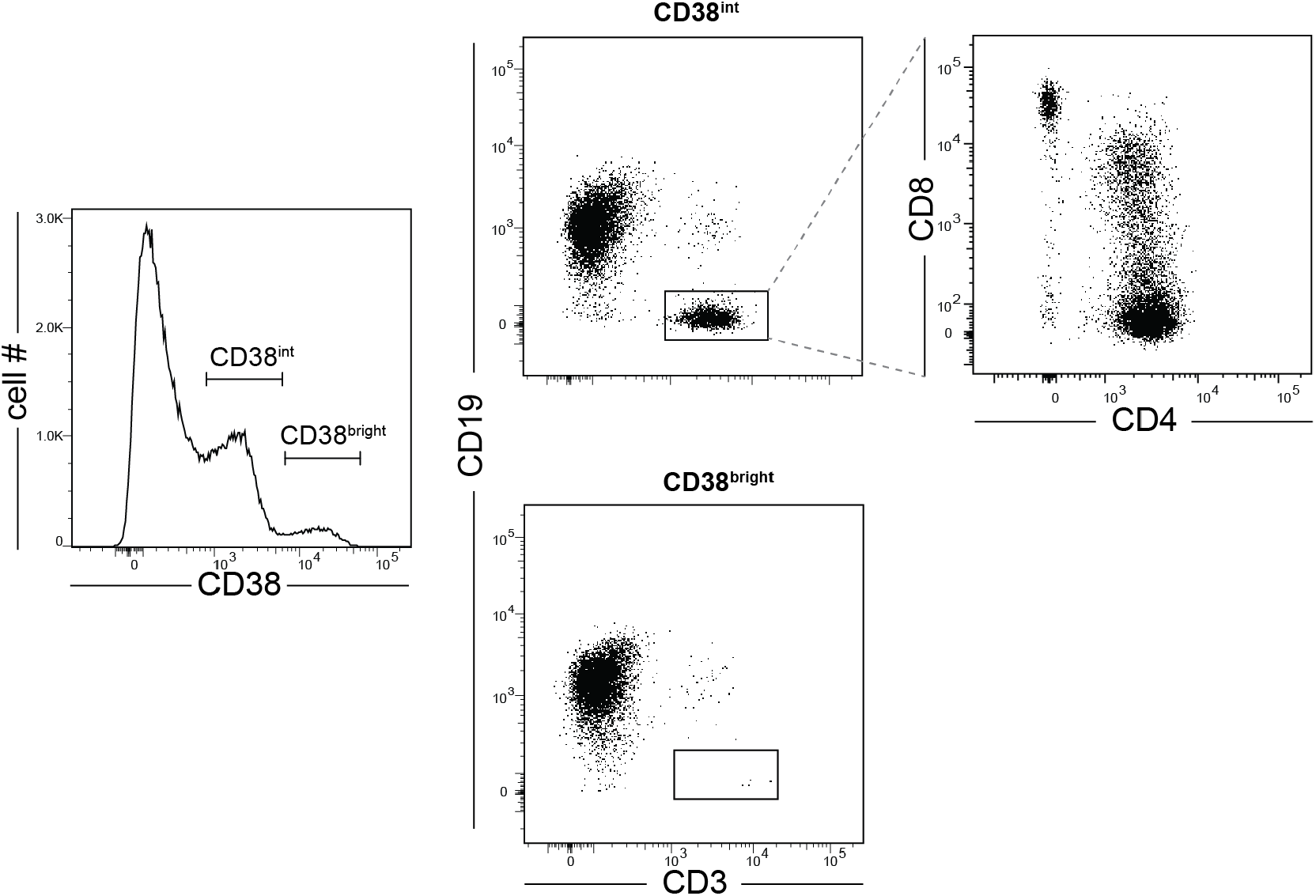
CD38 expression distribution across tonsillar mononuclear cells from a representative donor.

**Table S1.**
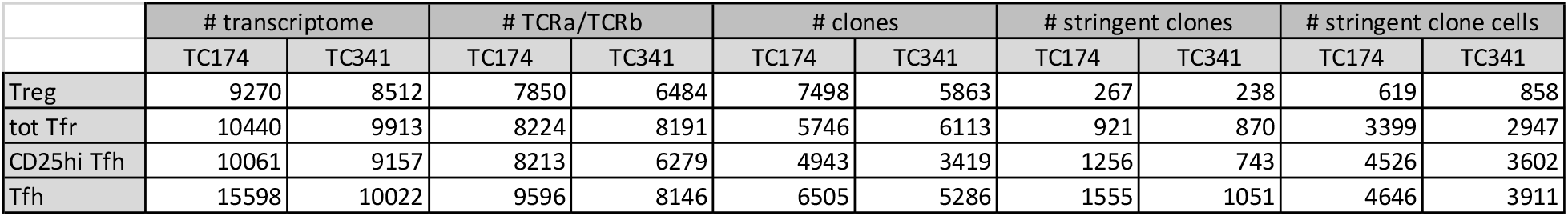
Summary statistics for single-cell experiments.

**Table S2.**
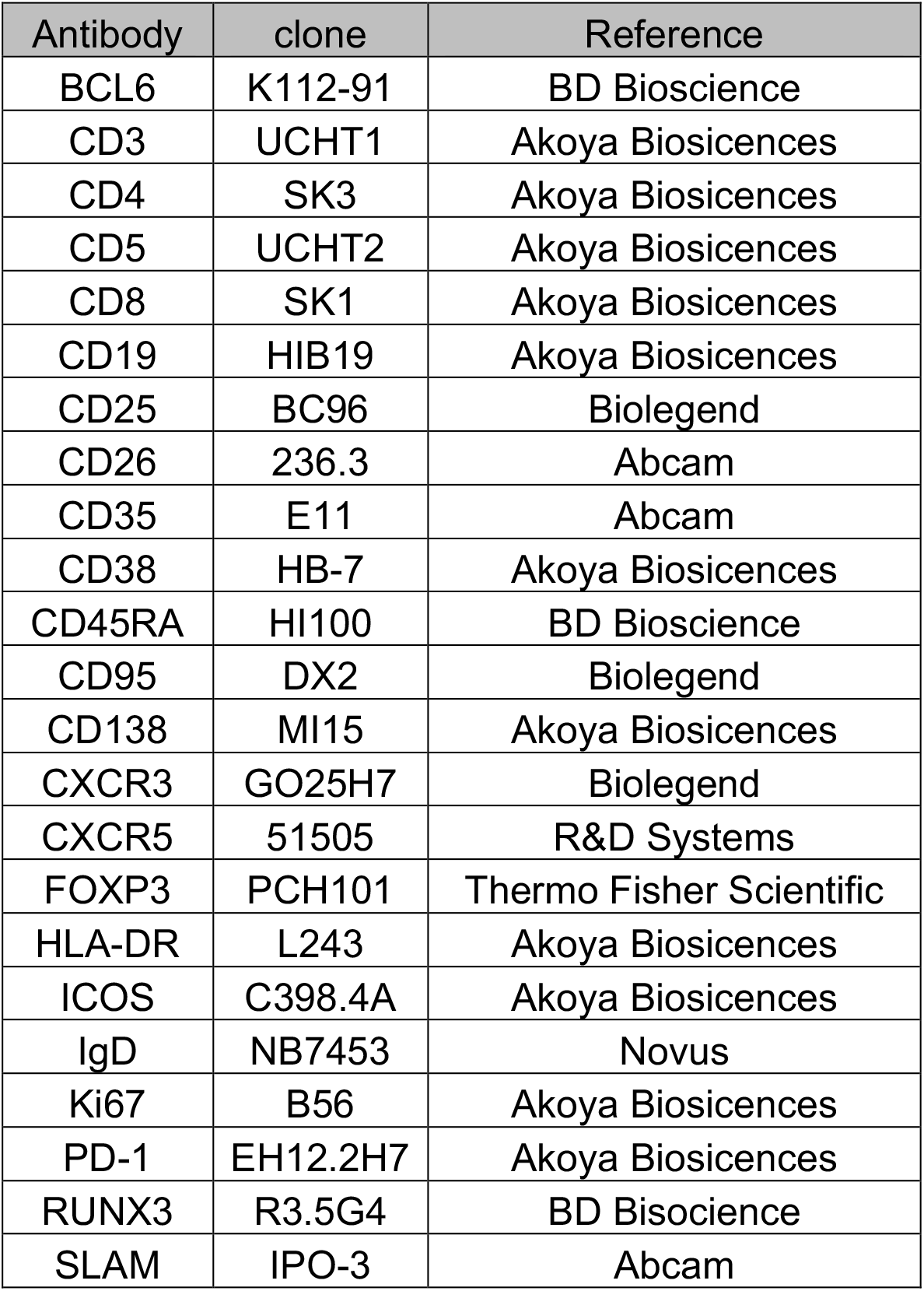
CODEX antibody panel.

