## Supplementary Materials for "Human T follicular helper clones seed the germinal center-resident regulatory pool"

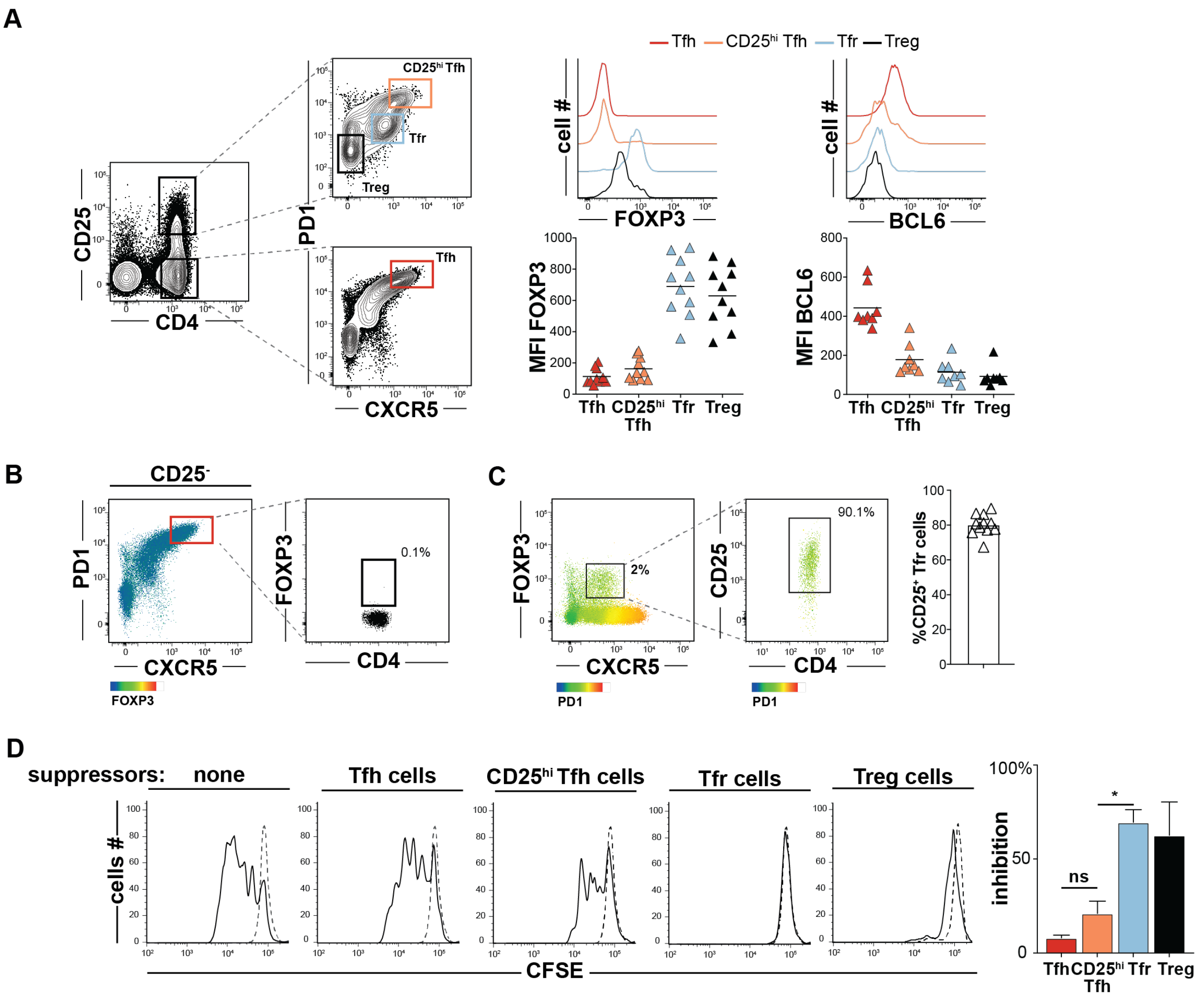


**Supplementary Figure 1. Treg and follicular T helper cell sorting strategy.** (**A**, left) Treg ($CD4^{+}CD25^{hi}CXCR5^{-}PD1^{-}$, black), Tfr **(**$CD4^{+}CD25^{hi}CXCR5^{+}PD1^{int}$, light blue), $CD25^{hi}$Tfh ($CD4^{+}CD25^{hi}CXCR5^{hi}PD1^{hi}$, orange) and Tfh ($CD4^{+}CD25^{-}CXCR5^{hi}PD1^{hi}$, red) gates are displayed on T cells from a representative tonsil donor. (**A**, right) Histograms and plots display FOXP3 and BCL6 mean fluorescence intensities (MFIs) on T helper subsets from a representative and all tonsil donors (n=6-8), respectively. Two-dimensional dot plots show (**B**) FOXP3 expression on cells in the Tfh gate and (**C**) PD1 expression on $FOXP3^{+}CXCR5^{+}CD25^{hi}$ Tfr cells from a representative donor and all donors (n=11). (**D,** left) Representative histograms of CFSE-labeled T-cell responders (Tresp, $CD4^{+}CD25^{-}CD45RO^{-}$) stimulated (solid line) or not (dashed line) in coculture with indicated tonsillar T helper cell subsets are displayed. Bar graph (right) represents mean percent inhibition relative to unstimulated Tresp cells from three tonsil donors. Error bars means ± SEMs. *, P<0.05 by Mann-Whitney U test.

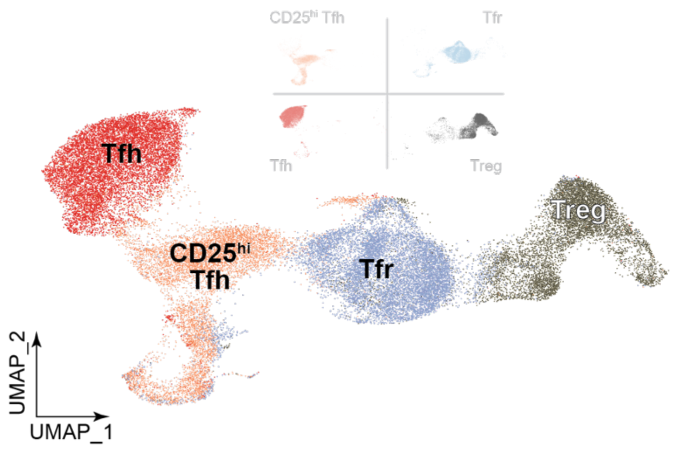


**Supplementary Figure 2. Dimensionally reduced transcriptomes of indicated TC341 cell subsets** are displayed as a UMAP.

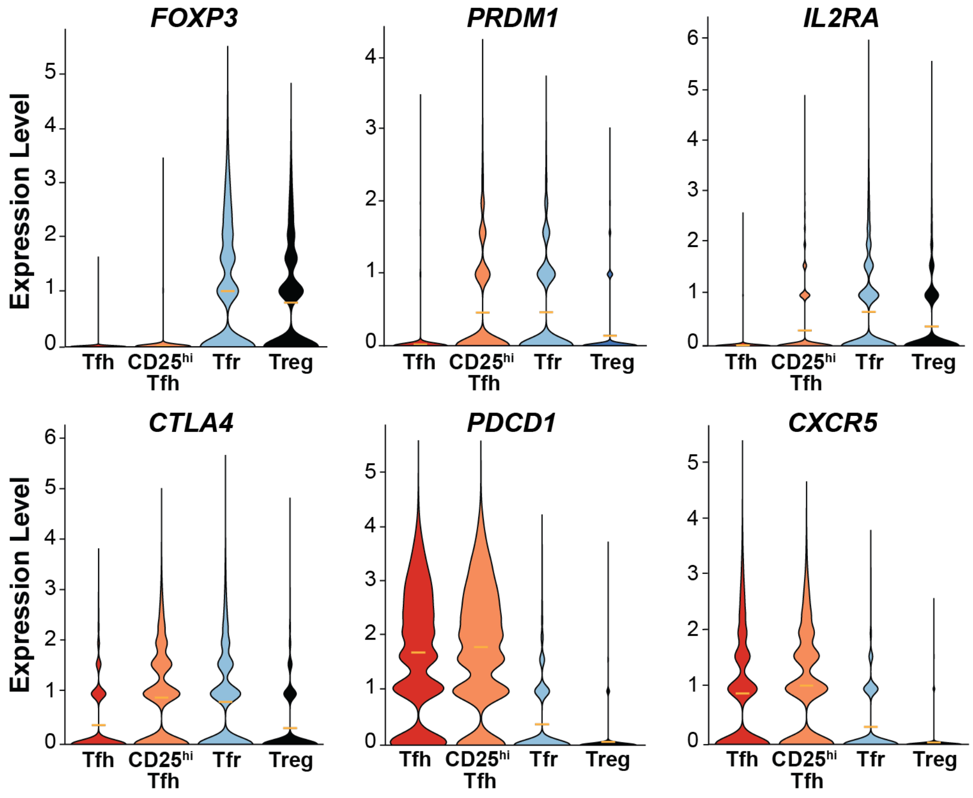


**Supplementary Figure 3. Violin plots display differentially expressed transcripts from pooled tonsil donor cells (TC174 and TC341).**

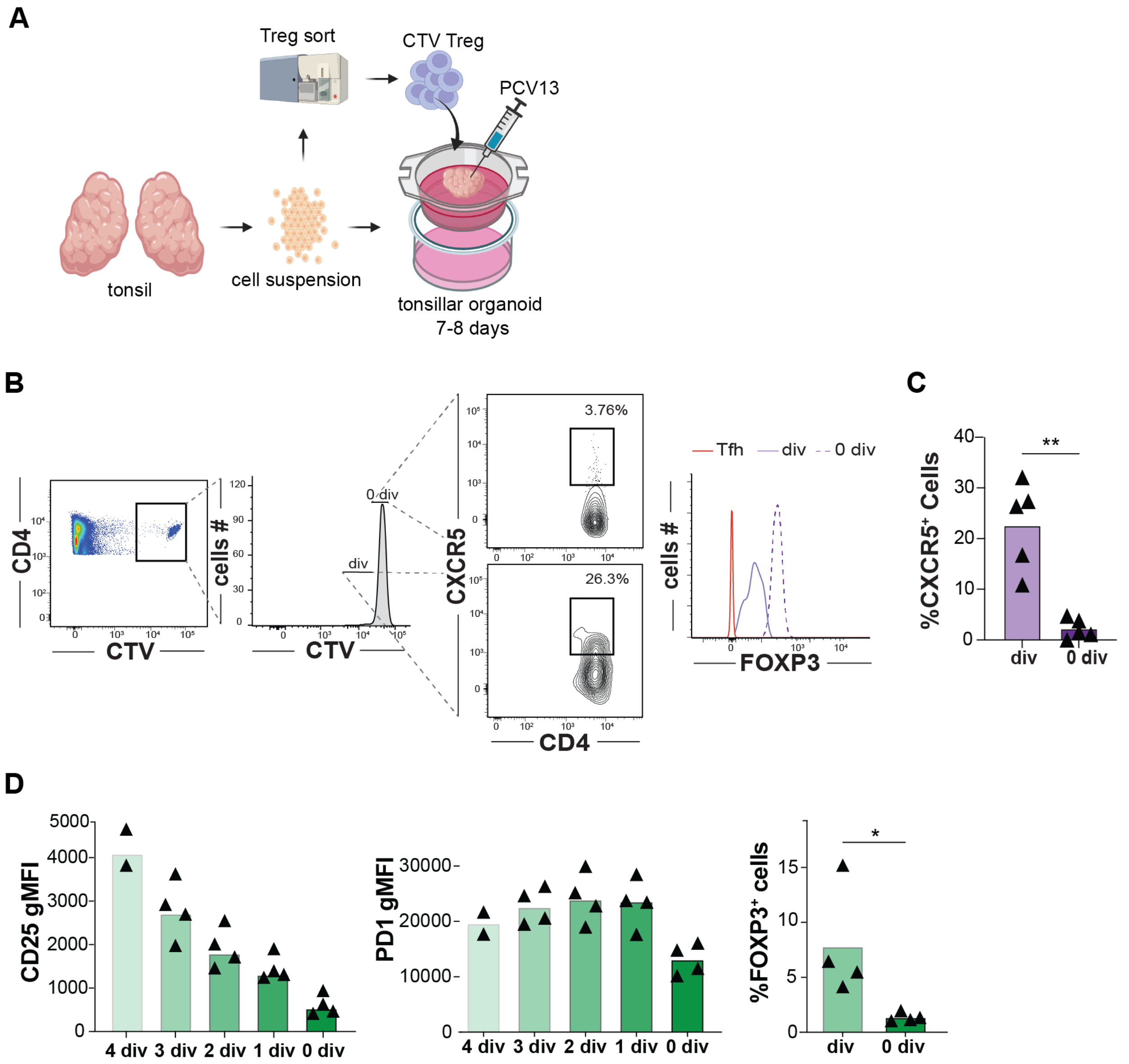


**Supplementary Figure 4. A strategy to track Tregs within vaccinated tonsillar organoids and measure differential protein expression on organoid incorporated Treg and Tfh cells.** (**A**) is depicted by cartoon. (**B**) Divided (div) Cell Trace Violet[TM] (CTV) stained Tregs from a representative organoid experiment differentially upregulate CXCR5 at day 7-8 compared with undivided (0 div) counterparts yet maintain FOXP3 expression. (**C**) CXCR5 expression of divided and undivided cells from five independent experiments are shown. (**D**) CD25, PD1, and FOXP3 expression by divided and non-divided CFSE stained-Tfh cells are displayed 7 days after PCV13 vaccination in tonsil organoids (n=4) *, $P<0.05$ ; **, $P<0.01$ by Mann-Whitney U test.

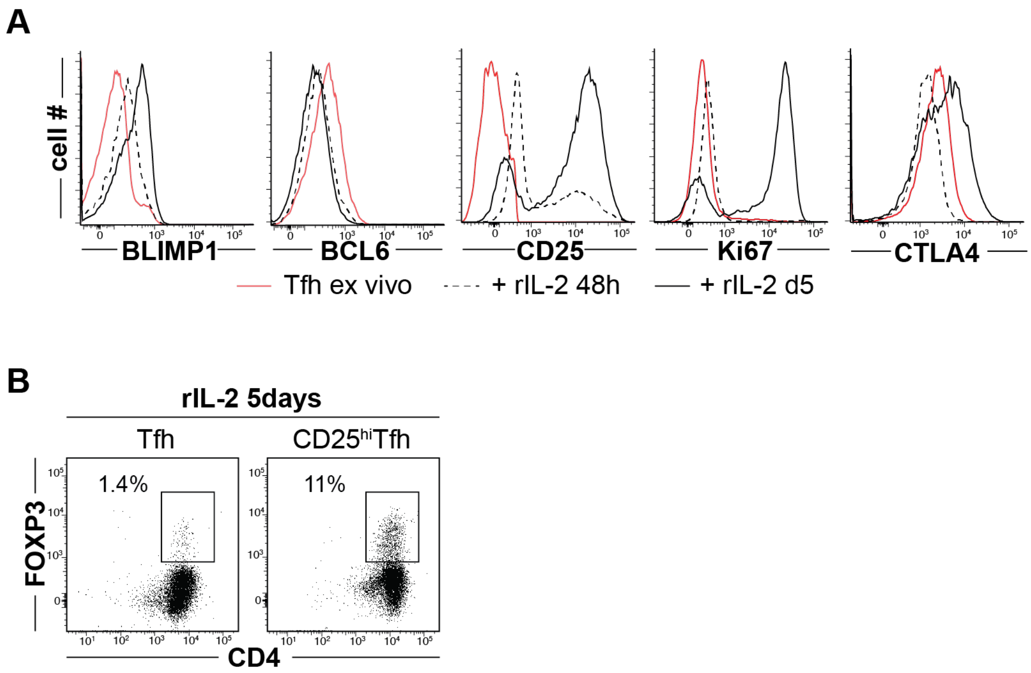


**Supplementary Figure 5.** ***In-vitro Tfh and CD25hiTfh cell differentiation experiments***. (**A**) Representative histograms of BLIMP1, BCL6, CD25, Ki67 and CTLA4 expression by Tfh cells before (red), after 48hours (dashed line) and five days (solid line) of recombinant IL2 treatment. (**B**) Representative dot plot of Tfh (left) and CD25$^{hi}$Tfh cell (right) FOXP3 expression after five days of recombinant IL2 (rIL-2) treatment.

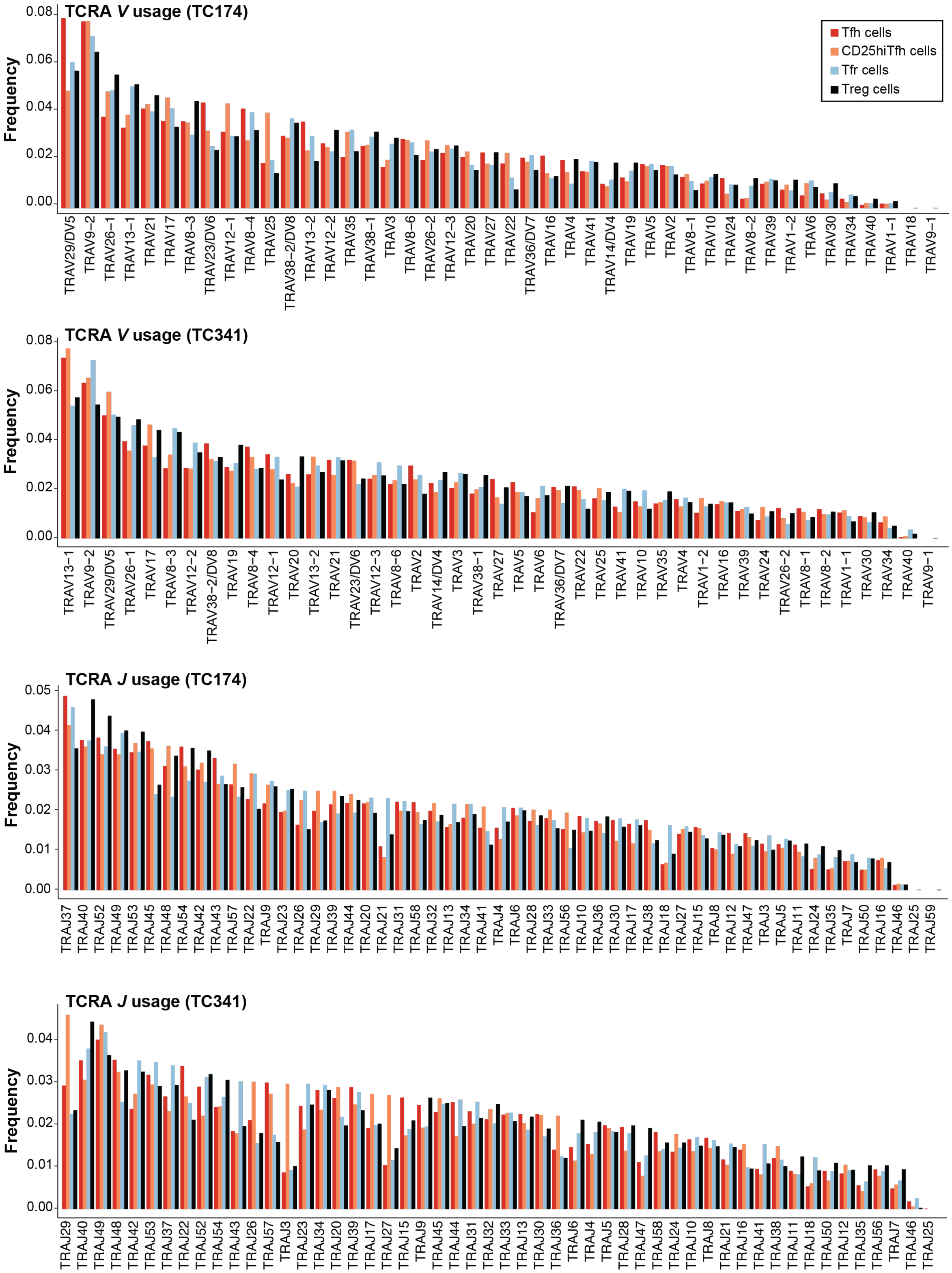


**Supplementary Figure 6. TCRA gene segment usage of TC174 and TC341.** Tfh cells (red), CD25$^{hi}$Tfh cells (orange), Tfr cells (light blue) and Tregs (black) TCR alpha chain *V* and *J* usage frequencies in tonsil donors TC174 and TC341.

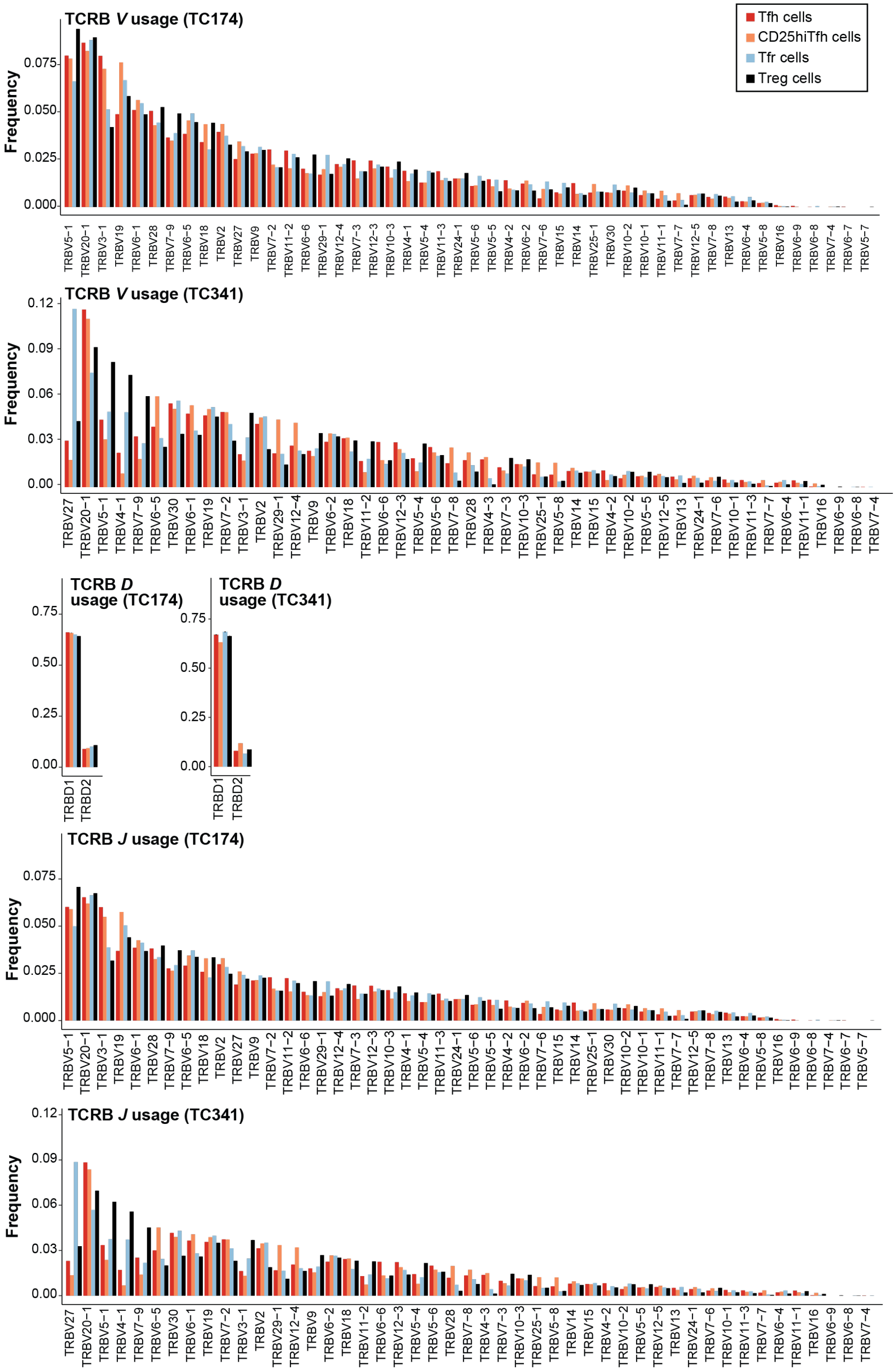


**Supplementary Figure 7. TCRB gene segment usage of TC174 and TC341.** Tfh cells (red), $CD25^{hi}$Tfh cells (orange), Tfr cells (light blue) and Tregs (black) TCR beta chain *V, D* and *J* gene segment usage frequencies in tonsil donors TC174 and TC341.

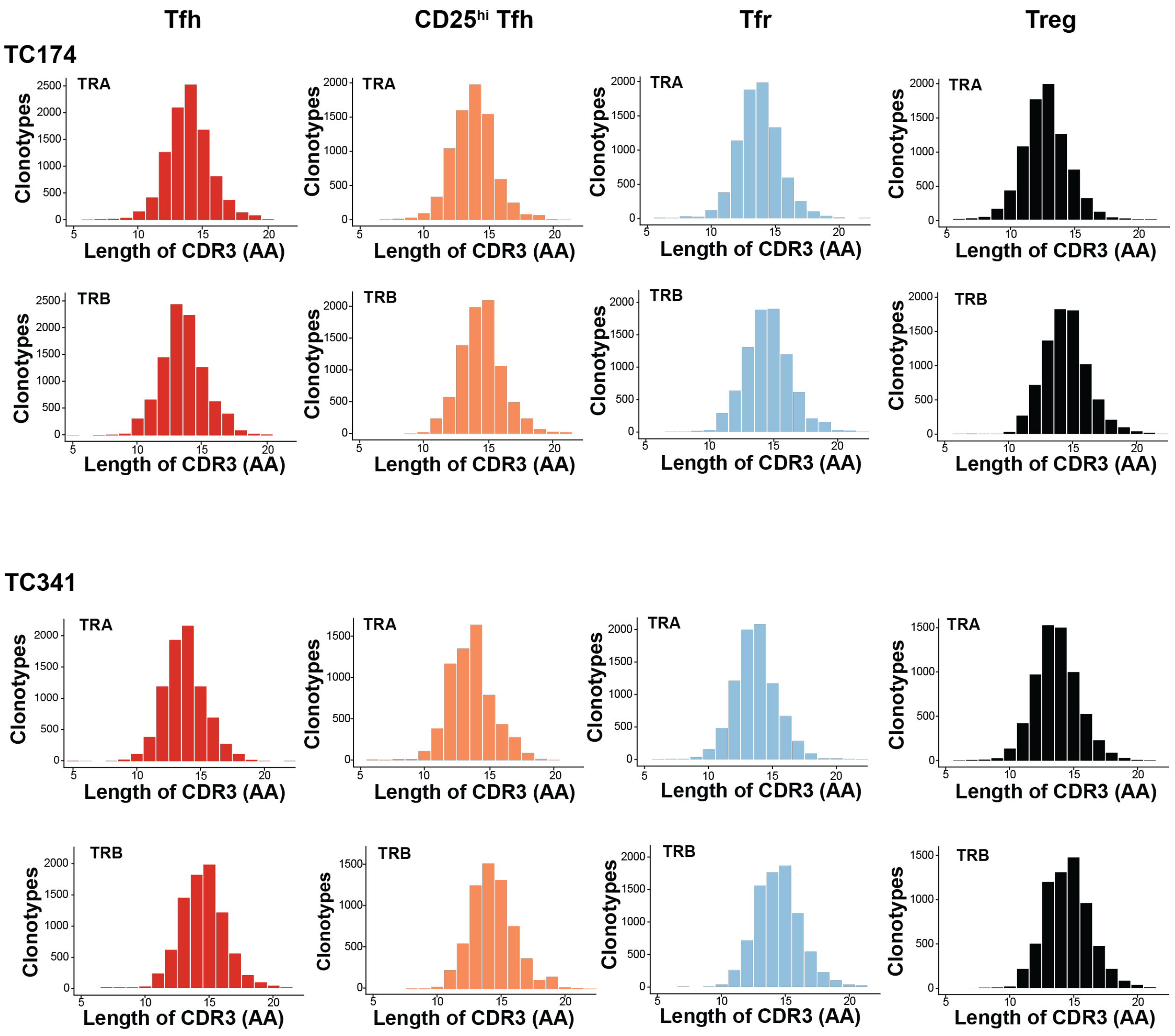


**Supplementary Figure 8. TCR CDR3 length distributions of TC174 and TC341.** Distribution of TCRA (above) and TCRB (below) CDR3 segments of indicated T-cell subsets (Tfh, red; CD25$^{hi}$Tfh, orange; Tfr, light blue and Treg, black) from two TC174 and TC341.

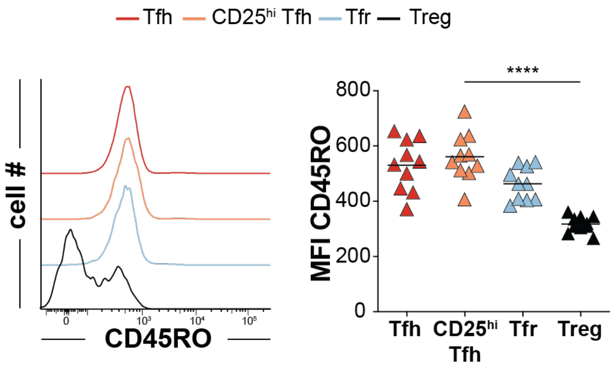


**Supplementary Figure 9. Differential CD45RO expression by CD4+ T cell subsets.** Tfh cells (red), CD25$^{hi}$Tfh cells (orange), Tfr cells (light blue) and Tregs (black) CD45RO are displayed as (left) histograms from a representative tonsil donor (right) and all tonsil donors (n=10). ****, *P*<0.0001 by Mann-Whitney U test.

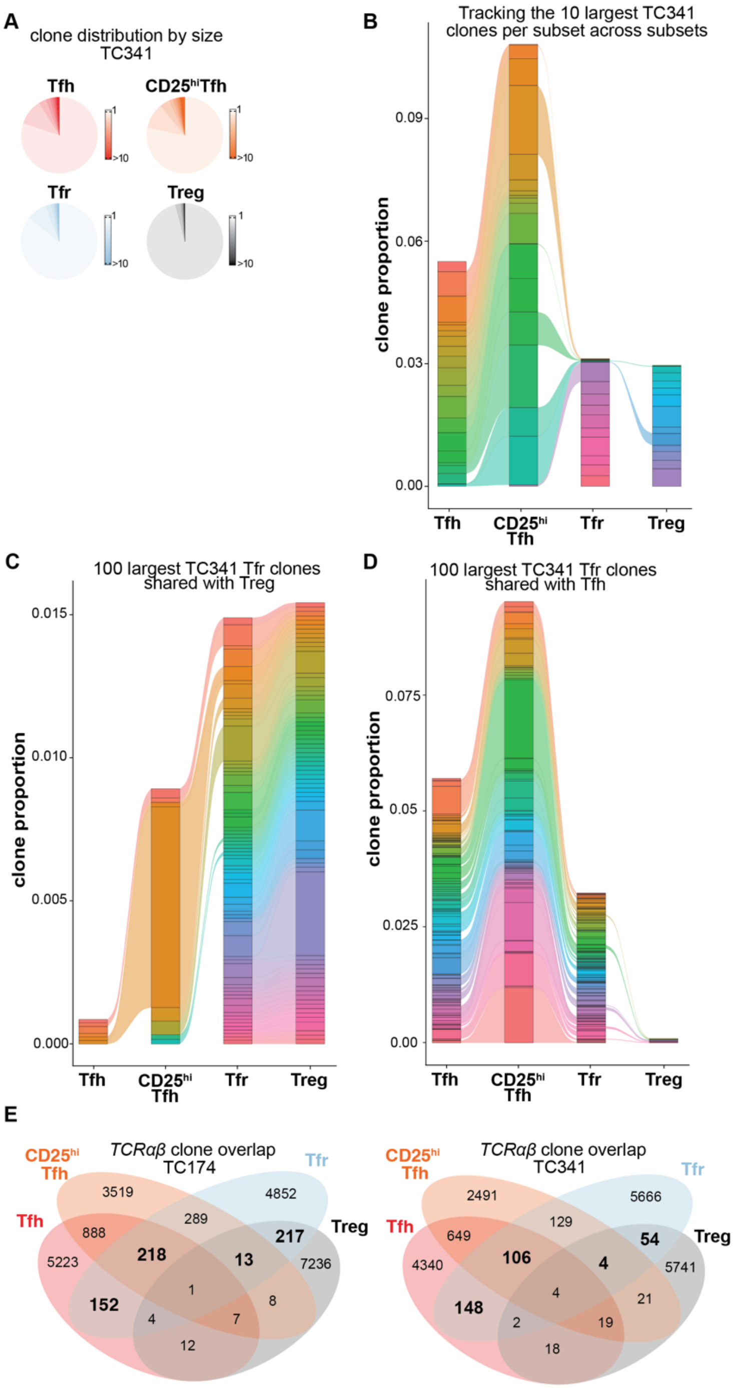


**Supplementary Figure 10. Clone size distribution and clone sharing between T helper subsets from TC174 and TC341.** (**A**) Pie charts indicate subset-specific clone size distributions (cells per clone) for TC341. (**B**) Tracking the top ten largest clones per subset across other subsets. (**C**) Tracking the top 100 largest clones shared between Tfr and Tfh cells and (**D**) between Tfr and Tregs across other subsets. More than ten clones or 100 clones per subset are sometimes displayed to account for ranking ties and clone sharing across subsets. (**E**) Venn diagrams depict clone sharing between T helper cell subsets from TC174 or TC341.

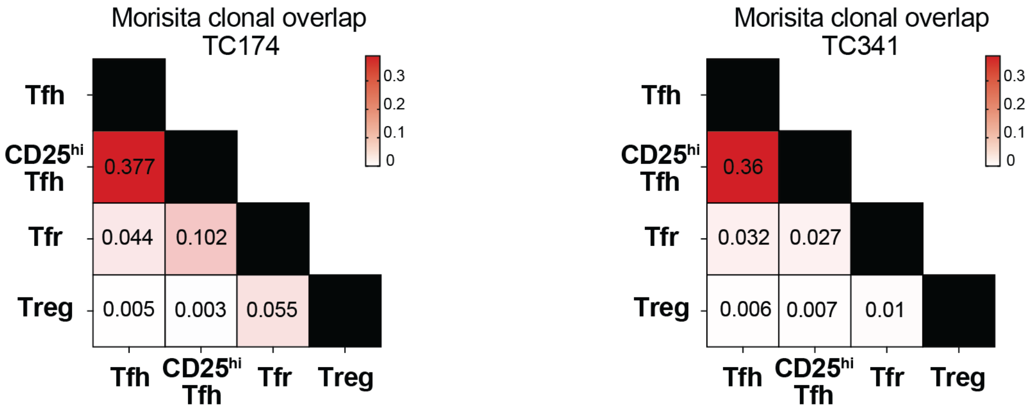


**Supplementary Figure 11. Morisita clonal overlap frequencies of TC174 and TC341.** T helper subsets from TC174 (left) and TC341 (right). Red shading indicates more clonal sharing.

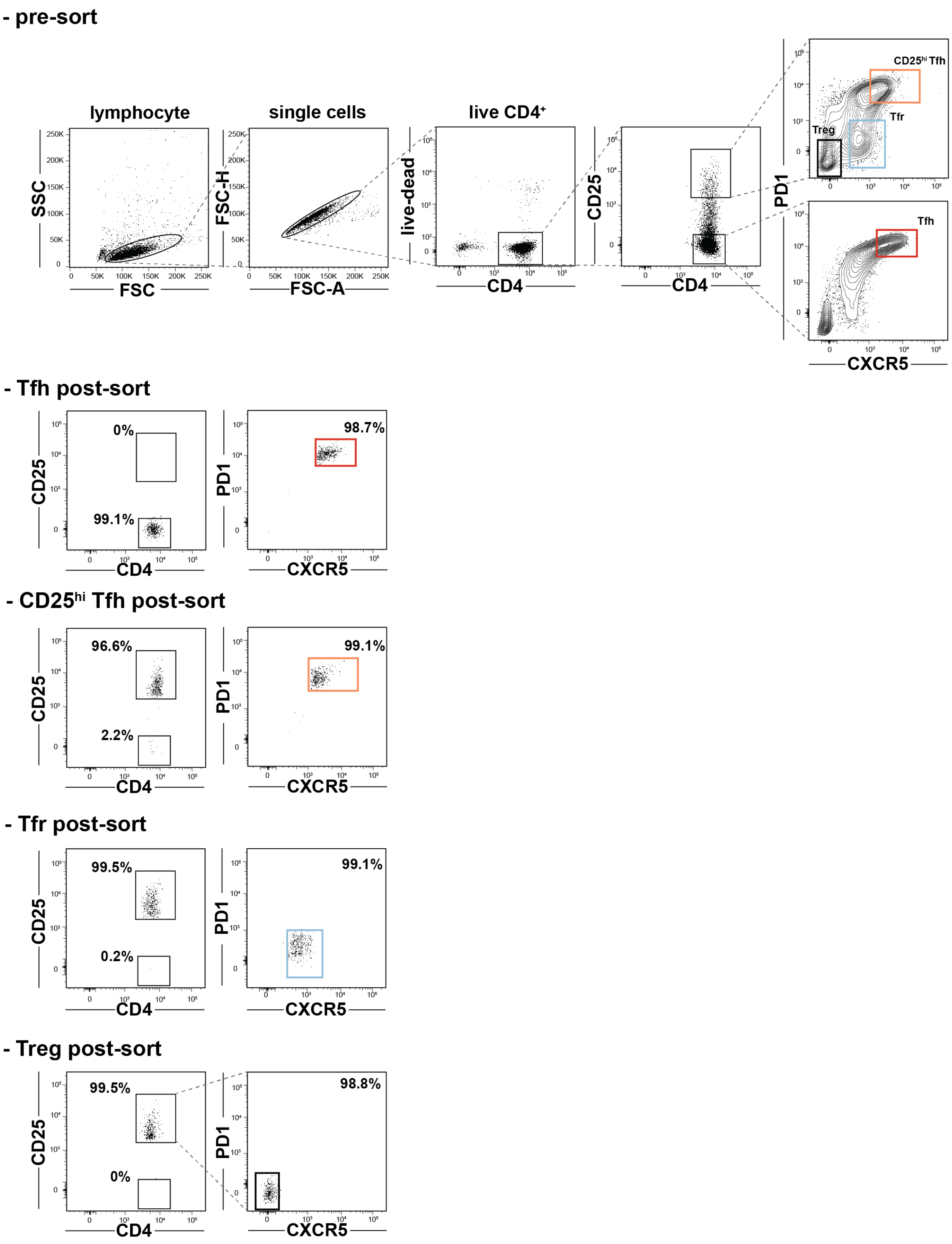


**Supplementary Figure 12. Pre and post-sort cell purity assessments.** (above) Sorting strategy to sort Tfh, CD25$^{hi}$Tfh, Tfr, and Treg cells. (below) Post-sort purities of the same subsets from a representative tonsil donor are displayed.

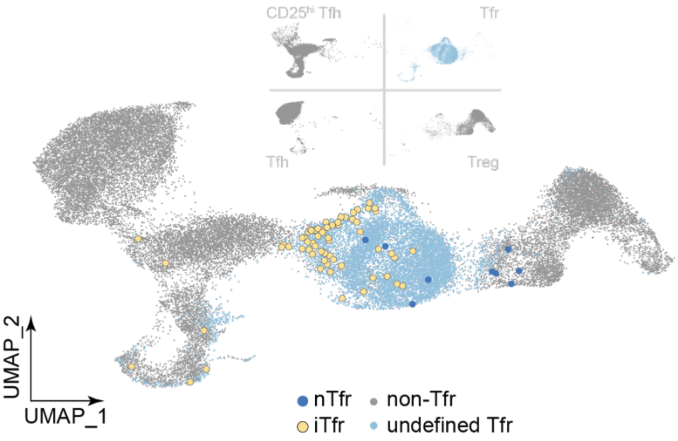


**Supplementary Figure 13. iTfr and nTfr localizations on the TC341 UMAP.** iTfr (yellow) and nTr (dark blue) cell positions are indicated relative to clonally-undefined Tfr cells (light blue) and non-Tfr cells (gray).

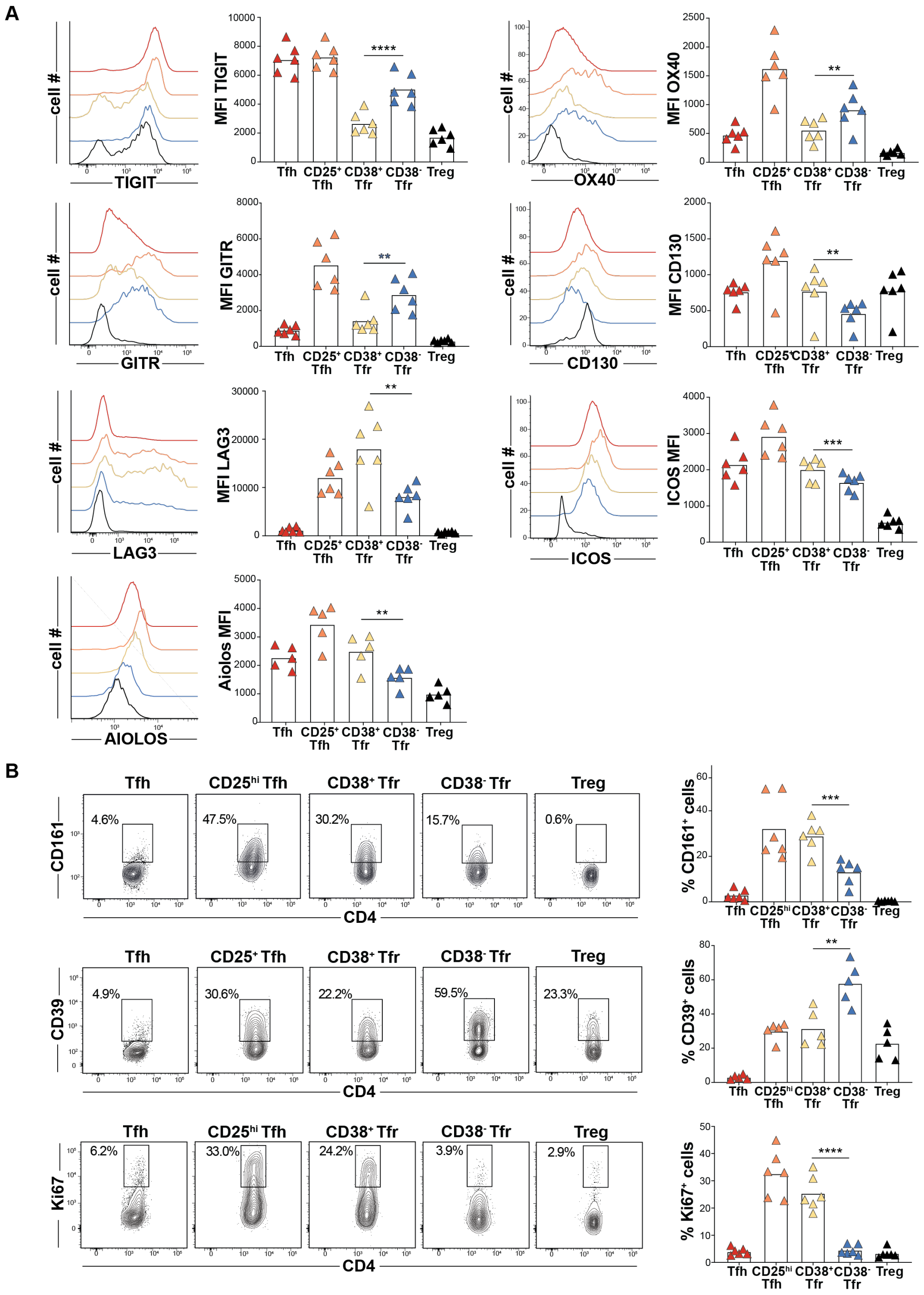

**Supplementary Figure 14. CD38⁺ and CD38⁻ Tfr cell extended immunophenotypes** (**A**) Histograms display TIGIT, OX40, GITR, CD130, LAG3 ICOS and AIOLOS expression by indicated T helper cell subsets from a representative tonsil donor (left) and bar graphs shows geometric mean fluorescence intensities (gMFIs) from counterpart cells fom five to six tonsil donors. (**B**) Dot plots present CD161, Ki67 and CD39 frequencies of indicated subsets from a representative tonsil donor (left) and five to six tonsil donors (right) **, *P*<0.01; ***, *P*<0.001; ****, *P*<0.0001 by Mann-Whitney U tests.

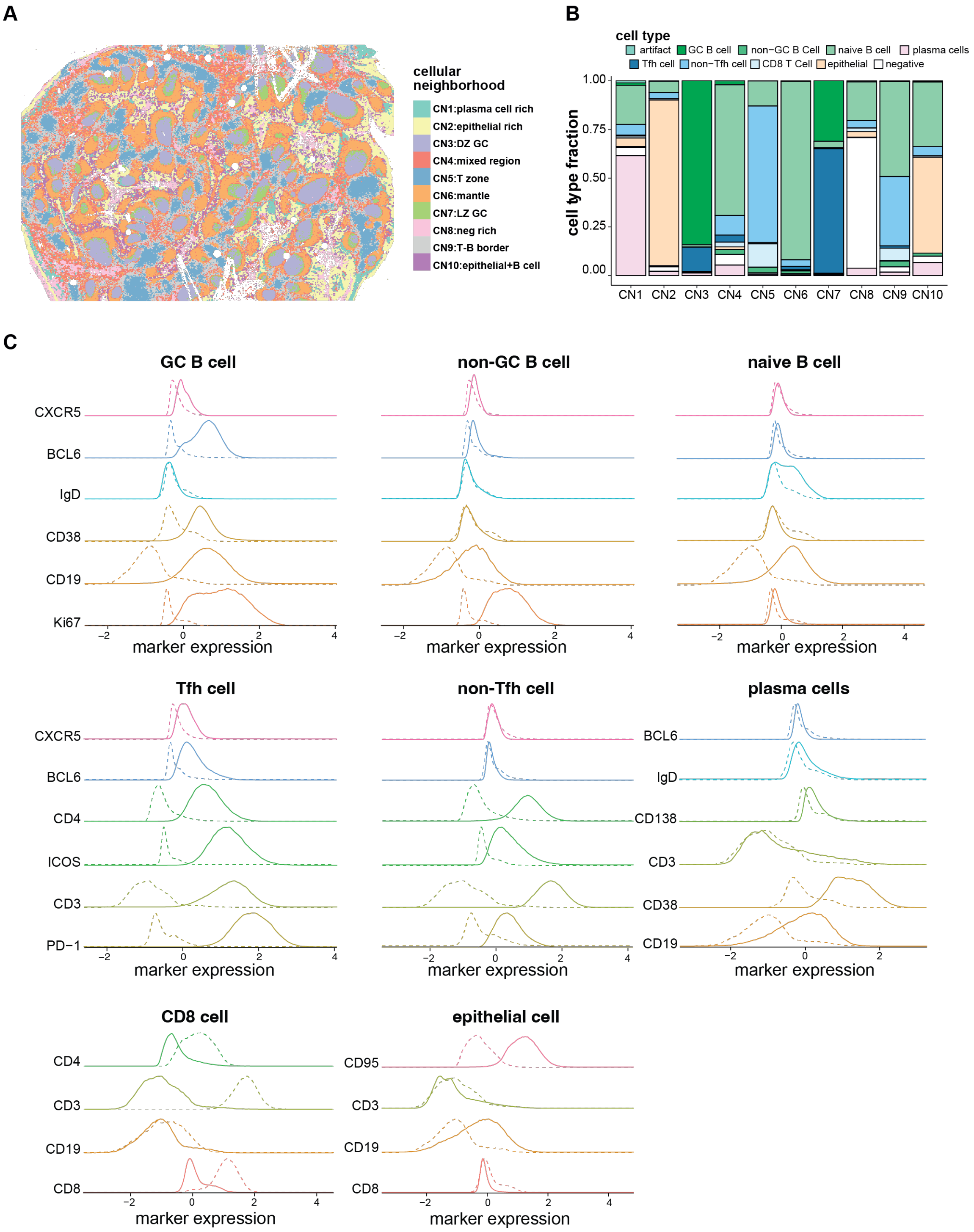


**Supplementary Figure 15**. **Tonsil cellular neighborhoods (CNs) and cell type profiles.** (**A**) CN positions, (**B**) cell type compositions by CN, and (**C**) cell type protein expression profiles of a CODEX-stained tonsil section are displayed.

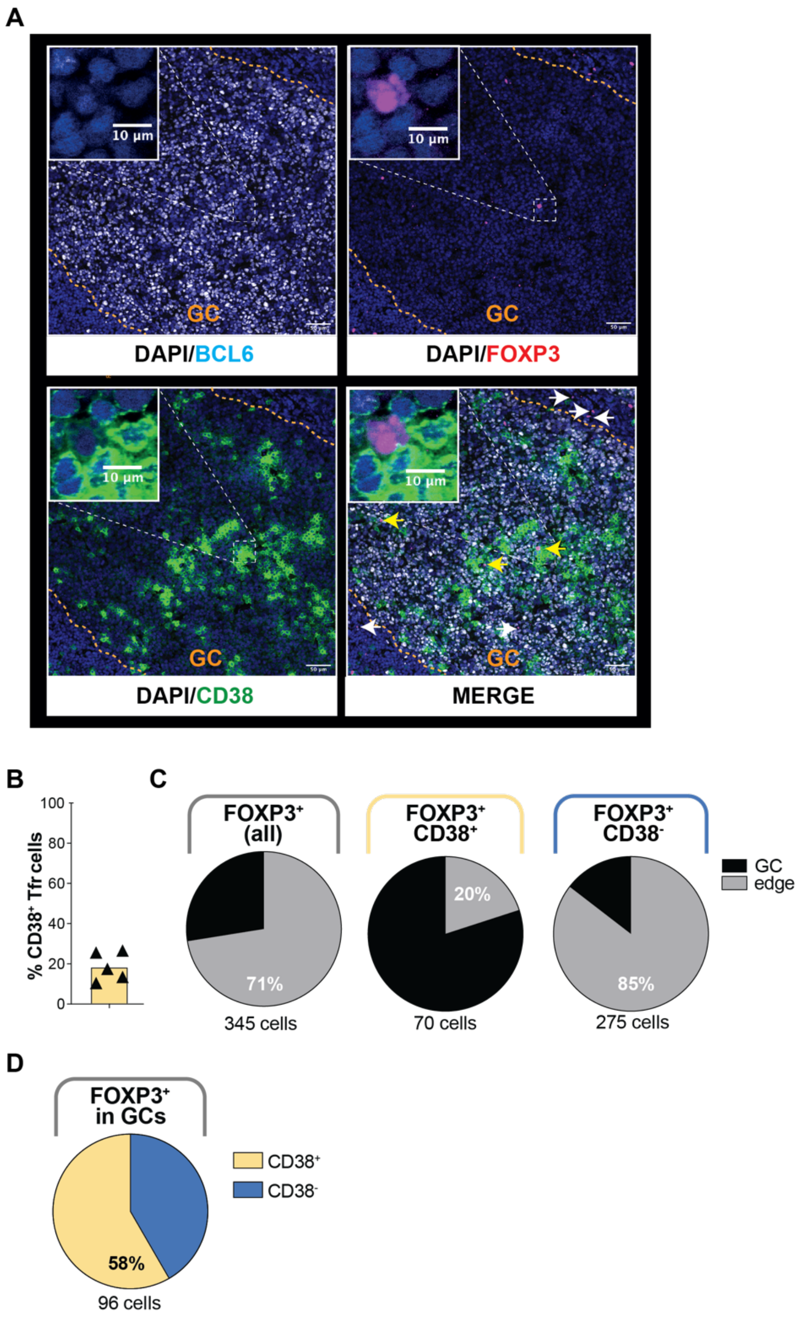


**Supplementary Figure 16. Confocal microscopy of stained tonsillar germinal centers (GCs).** (**A**) BCL6, FOXP3, CD38 and DAPI stains of a representative tonsil section are displayed separately and merged. The positions of $CD38^{+}FOXP3^{+}$ cells (yellow arrows) and $CD38^{-}FOXP3^{+}$ cells (white arrows) are indicated. White dashed squares contain one representative $CD38^{+}FOXP3^{+}$ cell magnified in insets at the top left corner. (**B**) Frequencies of GC-associated $FOXP3^{+}$ cells that express CD38, or not, from five tonsil donors are shown. (**C**) Proportions of $FOXP3^{+}$, $CD38^{-}FOXP3^{+}$, and $CD38^{+}FOXP3^{+}$ cells located within BCL6-defined GCs and outside them in the surrounding 50µm GC edge region and (**D**) Proportions of $CD38^{-}FOXP3^{+}$ and $CD38^{+}FOXP3^{+}$ cells located within BCL6-defined GCs are displayed.

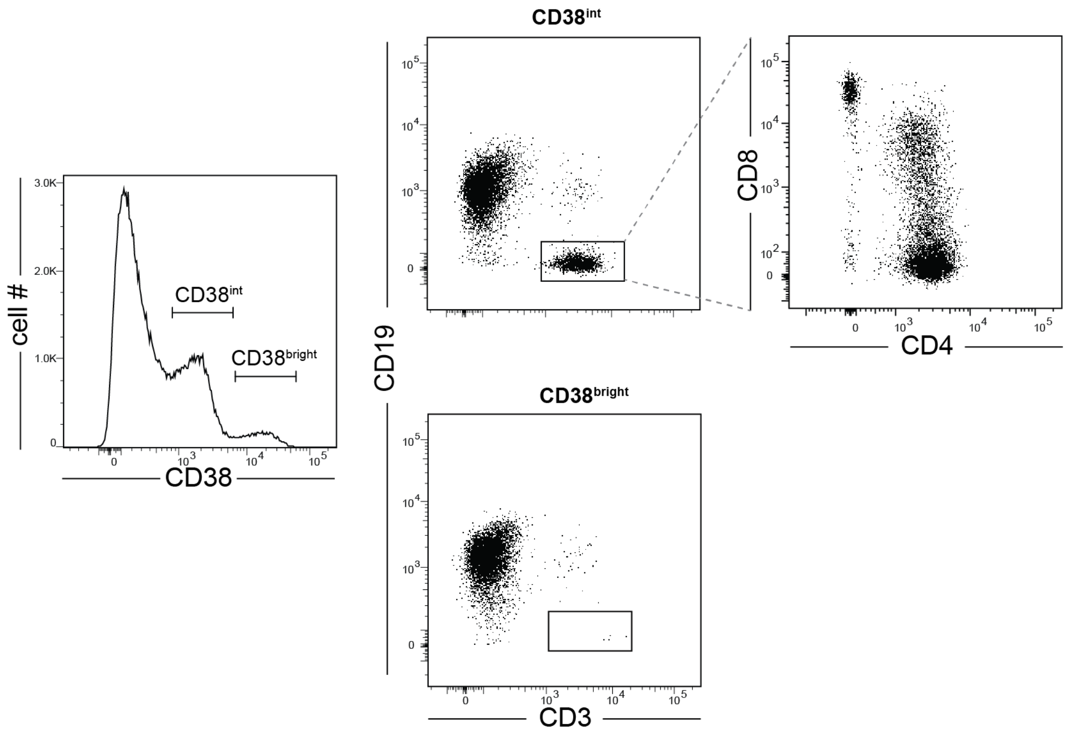


**Supplementary Figure 17. CD38 expression distribution across tonsillar mononuclear cells from a representative donor.**

**Table S1. Summary statistics for single-cell experiments.**

| | # transcriptome | | # TCRa/TCRb | | # clones | | # stringent clones | | # stringent clone cells | |
|---|---|---|---|---|---|---|---|---|---|---|
| | TC174 | TC341 | TC174 | TC341 | TC174 | TC341 | TC174 | TC341 | TC174 | TC341 |
| Treg | 9270 | 8512 | 7850 | 6484 | 7498 | 5863 | 267 | 238 | 619 | 858 |
| tot Tfr | 10440 | 9913 | 8224 | 8191 | 5746 | 6113 | 921 | 870 | 3399 | 2947 |
| CD25hi Tfh | 10061 | 9157 | 8213 | 6279 | 4943 | 3419 | 1256 | 743 | 4526 | 3602 |
| Tfh | 15598 | 10022 | 9596 | 8146 | 6505 | 5286 | 1555 | 1051 | 4646 | 3911 |

**Table S2. CODEX antibody panel**

| Antibody | clone | Reference |
|---|---|---|
| BCL6 | K112-91 | BD Bioscience |
| CD3 | UCHT1 | Akoya Biosicences |
| CD4 | SK3 | Akoya Biosicences |
| CD5 | UCHT2 | Akoya Biosicences |
| CD8 | SK1 | Akoya Biosicences |
| CD19 | HIB19 | Akoya Biosicences |
| CD25 | BC96 | Biolegend |
| CD26 | 236.3 | Abcam |
| CD35 | E11 | Abcam |
| CD38 | HB-7 | Akoya Biosicences |
| CD45RA | HI100 | BD Bioscience |
| CD95 | DX2 | Biolegend |
| CD138 | MI15 | Akoya Biosicences |
| CXCR3 | GO25H7 | Biolegend |
| CXCR5 | 51505 | R&D Systems |
| FOXP3 | PCH101 | Thermo Fisher Scientific |
| HLA-DR | L243 | Akoya Biosicences |
| ICOS | C398.4A | Akoya Biosicences |
| IgD | NB7453 | Novus |
| Ki67 | B56 | Akoya Biosicences |
| PD-1 | EH12.2H7 | Akoya Biosicences |
| RUNX3 | R3.5G4 | BD Bisocience |
| SLAM | IPO-3 | Abcam |
